# Evaluating key evidence and formulating regulatory alternatives regarding the UK’s Hunting Trophies (Import Prohibition) Bill

**DOI:** 10.1101/2023.06.13.544826

**Authors:** Dan Challender, Michael ’t Sas-Rolfes, Amy Dickman, Darragh Hare, Adam Hart, Michael Hoffmann, David Mallon, Roseline Mandisodza-Chikerema, Dilys Roe

## Abstract

Public policy addressing biodiversity loss is most likely to be effective when it is informed by appropriate evidence and considers potential unintended consequences. We evaluate key evidence relating to the Hunting Trophies (Import Prohibition) Bill that was discussed in the UK Parliament between 2022 and 2024. We characterize the UK’s role in international hunting trophy trade by analyzing CITES trade data for 2000-2021 and 2015-2021. For CITES-listed species imported to/exported from the UK as hunting trophies in these periods we use data from the International Union for Conservation of Nature Red List of Threatened Species to determine whether hunting designated as “trophy hunting” is i) likely a major threat contributing to species being of elevated conservation concern, ii) likely or possibly causing localized declines, or iii) not a threat. We then use the Red List to determine whether such hunting provides, or potentially provides, benefits for species and/or people. Finally, we evaluate the UK Government’s impact assessment of the bill. In 2000-2021 an estimated 3494 hunting trophies from 73 CITES-listed species and subspecies were exported to the UK involving an estimated 2549 whole organism equivalents (WOEs), i.e. individual animals. Imports involved 158.86 ± 66.53 (mean ± SD) trophies/year (115.83 ± 32.27 WOEs/year). In 2015-2021, 79% of imports were from countries where populations of the hunted species are stable, increasing, or abundant. Legal hunting for trophies is not a major threat to any of the species or subspecies imported to the UK, but likely or possibly represents a local threat to some populations of nine species. This hunting does, or could potentially, benefit 20 species and subspecies, and people. Among other concerns, the impact assessment failed to adequately consider the costs and benefits to local communities in countries where such hunting occurs. Informed by these analyses we discuss alternative regulatory options.

## 1. Introduction

Public policy addressing biodiversity loss is most likely to be effective when it is informed by appropriate evidence, is context-specific, and considers potential unintended consequences (Sutherland *et al*. 2020, IPBES 2022). Inadequate consideration of these factors can result in regulatory failure, including perverse impacts and counter-productive policies (Grabosky 1995, Baldwin *et al*. 2012). Overexploitation is a key threat to biodiversity (IPBES 2022) but devising policies to mitigate this threat can be inherently challenging. There is a lack of knowledge of many species, including their population biology, size, and trends, the impact of offtake, and the evolutionary impacts of harvesting (Smith *et al*. 2011). The most appropriate policies to address overexploitation may differ between contexts and scales related to ecological, economic, social, and/or governance factors (Cooney *et al*. 2015, IPBES 2022) meaning that identifying optimal solutions is not straightforward. Further complicating policy formulation regarding wildlife use are the frequently polarized (and sometimes misinformed) views of diverse stakeholders, especially concerning sentient and charismatic species, with ethical, ideological, and scientific arguments used to support or oppose potential options (Hammond *et al*. 2022, Mkono 2022). Yet with adequate knowledge of species and the social-ecological systems that they are part of, appropriate policies can be devised to support species conservation and benefit local people (’t Sas-Rolfes *et al*. 2022).

“Trophy hunting” – defined by the IUCN (2012) as legal, low-offtake hunting, where hunters pay a high fee to hunt individual animals with particular characteristics (e.g., horn length) and retain all or part of the animal – has emerged as a contemporary conservation policy debate, with widespread media coverage, particularly following the death of Cecil the lion in 2015 (’t Sas-Rolfes 2017, Yeomans *et al*. 2022). This has largely been due to advocacy groups arguing that such hunting threatens wildlife populations, disregards animal welfare, is morally reprehensible and should be further legislated against (Born Free *et al*. 2022). Conversely, evidence indicates that in diverse circumstances across several continents, legal and well-managed hunting for trophies can deliver benefits to local people and support conservation by ensuring that biodiversity is a competitive land use option (IUCN 2016a, Parker *et al*. 2023). Nevertheless, various governments have legislated to restrict the trade in hunting trophies since 2015. For example, Australia and France have banned imports of lion (*Panthera leo*) trophies, Finland has banned imports of trophies from all species on Annex A and selected species on Annex B of the EU Wildlife Trade Regulations (EUWTRs), and the Netherlands has prohibited trophy imports from over 200 species (Ares 2019).

In the last decade, advocacy groups have encouraged the United Kingdom (UK) to tighten controls on the import and export of hunting trophies (Ares 2019). The Wildlife Trade Regulations (WTRs) in the UK (as retained EU law) and EUWTRs in Northern Ireland list all CITES (Convention on International Trade in Endangered Species of Wild Fauna and Flora) and some non-CITES species in four Annexes (A–D). Under these regulations, imports of hunting trophies from wild species to the UK require an import and export permit for Annex A species and six species in Annex B: white rhino (*Ceratotherium simum*), hippopotamus (*Hippopotamus amphibius*), African elephant (*Loxodonta africana*), argali (*Ovis ammon*), polar bear (*Ursus maritimus*), and lion. For other species in Annex B only an export permit is required, and hunting trophies are treated as personal and household effects. Import and export permits can only be granted based on non-detriment findings (NDFs), to ensure that trade is not detrimental to wild populations. International trade in hunting trophies involving CITES-listed species is therefore regulated with exporting countries providing key oversight.

In 2019 the UK Government issued a call for evidence on the scale of hunting trophy imports and exports and associated impacts and held a consultation on further restricting this trade. In June 2022, a Private Members’ Bill – the Hunting Trophies (Import Prohibition) Bill – proposed to ban the import of hunting trophies to the UK from species listed in Annexes A and B of the WTRs, which the Government stated would protect ∼7000 species (UK Government 2021a). The rationale was to ensure that imports of hunting trophies to the UK do not place additional pressure on species of conservation concern and, based on the belief that since the British public feels strongly about “trophy hunting”, it is an issue the government should address (DEFRA 2021). The bill failed to pass the second committee stage in the House of Lords in September 2023. An identical Private Members’ Bill was submitted in December 2023 but did not progress to the committee stage in the House of Commons. This article pertains to both bills and to future legislation with similar intentions.

The rationale for an import ban asserts that legal hunting for trophies threatens many species, including those imported to the UK, but the evidence to support this is unclear. Here, we evaluate key evidence relating to the bill and the associated policymaking process. We focus on CITES-listed species because these have been deemed in need of international trade regulation to avoid overexploitation (CITES 1973). Import and export data are routinely collected for CITES-listed species in the UK but not for other species.

Specifically, we:

1) estimate the number of hunting trophies from CITES-listed species and the associated number of animals traded globally in the periods 2000-2021 and 2015-2021.
2) characterize the role of the UK in this trade considering the species involved, the number of trophies and individual animals traded, the source of trophies (e.g., wild vs. captive-bred), and importing and exporting countries. We focus on the period 2000-2021 to provide historical context and the period 2015-2021, which broadly aligns with the UK Government’s impact assessment (see DEFRA 2021) (Supp. Material 1).
3) determine the population status of CITES-listed species exported to the UK as hunting trophies in the period 2015-2021 and calculate the proportion of trade sourced from populations with different status.
4) contextualize UK hunting trophy imports of CITES-listed species in overall UK trade in animal species listed under CITES and trade for commercial purposes and as pets.
5) use data from the International Union for Conservation of Nature (IUCN) Red List of Threatened Species (hereafter “Red List”) to determine for CITES-listed species imported to/exported from the UK whether legal hunting for trophies (as defined) is i) likely a major threat contributing to species being of elevated conservation concern, 1. ii) either likely or possibly causing localized declines (distinction explained below), or iii) not a threat, and additionally whether it provides, or has the potential to provide, benefits for species and/or local communities where such hunting takes place.
65) use quality indicators from the Regulatory Policy Committee (RPC; Regulatory Policy Committee, n.d.) to evaluate the UK Government’s impact assessment of the Hunting Trophies (Import Prohibition) Bill in conservation terms.

Informed by these analyses, we discuss alternative policy options that UK policymakers may wish to consider in regulating international trade in hunting trophies of CITES-listed species. This article should be of interest to politicians and policymakers in the UK, the EU and other countries considering similar legislation, countries where trophy hunting takes place, and practitioners and academics in the UK and internationally.

## 2. Methods

### 2.1 International trade in CITES-listed species

To estimate the number of hunting trophies from CITES-listed species and associated number of animals traded globally, characterize the role of the UK in this trade, and contextualize UK hunting trophy imports within overall UK trade in CITES-listed animal species and compare these imports with trade for commercial purposes and as pets, we used CITES trade data. In March-May 2023 we downloaded comparative tabulation reports from the CITES trade database for the period 2000-2021 (CITES Trade Database 2023) recognizing that the latest year of complete data is expected to be 2021, i.e., two years prior to the current year (CITES Secretariat and UNEP-WCMC 2022). For search terms used see Supp. Material 1. We summarized direct trade using pivot tables in MS Excel and used exporter-reported quantities because they are often more complete (CITES Secretariat and UNEP-WCMC 2022), recognizing that these quantities may refer to permits issued rather than quantities of specimens exported (Supp. Material 1). For trade terms and units used see Supp. Material 1. As one animal may produce multiple trophies, the number of trophies does not equate to number of animals killed but converting trade volumes to whole organism equivalents (WOEs) enables estimates of the number of individual animals involved (Supp. Material 1). We estimated WOEs for trade adapting the approach by Harfoot *et al*. (2018) (Supp. Material 1). We used RStudio Version 1.4.1717 to calculate means and standard deviations for trade in species and WOEs over time.

### 2.2 Population status

We determined the population status (e.g., increasing, decreasing or stable) of each species exported to the UK as a hunting trophy in the period 2015-2021 by reviewing and collating the available information for each exporting country in the global Red List assessment for each taxon (using Red List version 2022-2, IUCN 2023) and/or regional assessments where they were available. We then calculated the proportion of trade sourced from countries with populations of these species that are increasing or decreasing or similar respectively.

### 2.3 Trophy hunting as a contributor to species being of elevated conservation concern and as a benefit provider

To determine whether legal hunting for trophies is i) likely a major threat contributing to species being of elevated conservation concern (i.e., to a species meeting, or approximating, the thresholds for listing in any of categories NT [Near Threatened], VU [Vulnerable], EN [Endangered], CR [Critically Endangered] or EW [Extinct in the Wild], as defined by IUCN), ii) either likely or possibly causing localized declines, or iii) not a threat to species, we built a MS Excel database including data from Red List assessments (using version 2022-2) for species imported to/exported from the UK as hunting trophies in 2000-2021 (Supp. Material 1). Our approach to classifying species in the above three categories draws on a similar analysis published for international trade (Challender *et al*. 2023). We read the narrative text in the threats and justification fields of each assessment and interpreted this information with available information on coded threats for species (e.g., 5.1.1. [Hunting & collecting terrestrial animals → Intentional use [species being assessed is the target]]) (Supp. Material 1). We considered the timing (e.g., past), scope (e.g., minority of the population), and severity (e.g., causing rapid declines) of the threats (where these data were available), to assist in distinguishing between major and minor threats (IUCN 2016b). We read the Use and Trade field of assessments for additional context and used information on how species are used and/or traded to inform decision-making (Supp. Material 1).

We categorised legal hunting for trophies as a major threat contributing to species being of elevated conservation concern where the available evidence indicates that a species has intentional use as a major threat (based on the threats narrative and/or coded threats, including timing, scope, and severity), and the threats narrative indicates that trophy hunting (as defined) is currently a primary factor driving this threat. Where this was not the case, species were not considered to have legal hunting for trophies as a major threat. We then categorised these species into two sub-categories based on whether there is evidence indicating that legal hunting for trophies is either a) *likely* (i.e., probable) or b) *possibly* (i.e., stated but qualified as uncertain, e.g., “may be”, “potentially” or similar) causing localized declines. Species that did not meet either of these criteria were considered not to be threatened by legal hunting for trophies at any level. We determined, based on available information in Red List assessments, whether legal hunting for trophies does, or has the potential to, benefit the species and/or local people (Supp. Material 1). The Red List was updated in December 2023, and we checked and updated our categorization of species against Red List version 2023-1 (IUCN 2024).

We acknowledge that there are limitations to using Red List data to document whether species are used and/or traded, and whether species are threatened by use and/or trade. These limitations, discussed elsewhere (Challender *et al*. 2022, 2023; Marsh *et al*. 2022), include that assessments need updating and may omit information on use and/or trade of species. There may also be biases in taxonomic groups assessed on the Red List, but most species hunted for trophies have been assessed meaning we can be confident our approach is robust.

### 2.4 Evaluating the UK Government’s impact assessment of the bill

We used quality indicators from the RPC to evaluate the UK Government’s impact assessment of the bill in conservation terms (Supp. Material 1). Regulatory proposals in the UK must be accompanied by an impact assessment which evaluates the likely risks, costs, and benefits of proposed regulation to businesses, the public and third sectors, and individuals in the UK (Regulatory Policy Committee, n.d.). Guidance states that it is sometimes reasonable to consider people living outside of the UK, and that impacts on wildlife and the natural environment should be considered (Supp Material 1).

The RPC is an independent regulatory body, which assesses the quality of evidence and analysis used to inform UK Government regulatory proposals (Regulatory Policy Committee, n.d.). We used the RPC’s quality indicators to evaluate the impact assessment for the bill (DEFRA 2021), considering whether assumptions were reasonable and justified, the quality of analysis and evidence, and areas of the assessment that could be improved (Supp. Material 1). We applied this to all five sections of the impact assessment: policy rationale, costs and benefits, risks and unintended consequences, wider impacts, and post implementation review.

## 3. Results

### 3.1 International trade in hunting trophies of CITES-listed species, the role of the UK, and population status of imported species

Direct trade in hunting trophies of CITES-listed species in 2000-2021 (i.e., over 22 years) involved an estimated 557,799 trophies globally and an estimated 419,877 WOEs. The top exporters were Canada and South Africa (>130,000 trophies respectively) while the top importer was the US, importing >247,000 trophies (Supp. Material 2). The UK ranked 19/74 exporting countries, exporting an estimated 968 trophies (968 WOEs). Among importing countries, the UK ranked 25/183, importing an estimated 3494 trophies (2549 WOEs), <1% of the global trade in terms of number of trophies and WOEs.

In 2015-2021 (i.e., 7 years) trade globally involved an estimated 162,891 hunting trophies from CITES-listed species (106,005 WOEs). The top exporters were South Africa (53,304 trophies) and Canada (36,520) while the top importer was the US (74,107 trophies) (Supp. Material 2). The UK ranked 37/53 exporters, exporting 17 trophies (17 WOEs), and 25/155 importers, importing an estimated 951 trophies (738 WOEs), again <1% of global trade.

The UK imported hunting trophies from 73 CITES-listed species and subspecies in 2000-2021, mainly involving mammals (96%) (Supp. Material 2). The top 10 imported species accounted for 74% of this trade (2569/3494 trophies), which mainly involved wild specimens (Table 1). Overall, this trade involved an estimated 158.86 ± 66.53 (mean ± SD) trophies/year (115.83 ± 32.27 WOEs/year). For African elephant, the species with the highest number of trophy imports, this involved 24.45 ± 30.23 trophies/year, or 5.66 ± 5.08 WOEs/year. Most (88%) trade in the top 10 imported species came from just six countries: South Africa (29%), Canada (18%), Zimbabwe (13%), Namibia (13%), Botswana (8%), and Zambia (7%). For 63 species and subspecies (86%), a mean of <5 trophies were imported annually (Supp. Material 2). Of the 73 species and subspecies, 50 (69%) averaged imports of less than one trophy a year, and 17 (23%) had a single trophy imported over the 22-year period (Supp. Material 2).

**Table 1.** Top ten species imported to the UK as hunting trophies in 2000-2021 ranked by number of trophies, mean no. of trophies imported/year, WOEs imported, and mean WOEs imported/year. . Source: CITES Trade Database (2023). WOEs=Whole organism equivalent. BW=Botswana, CA=Canada, CM=Cameroon, MW=Malawi, MZ=Mozambique, NA=Namibia, TZ=Tanzania, UG=Uganda, US=United States of America, ZA=South Africa, ZM=Zambia, ZW=Zimbabwe. CITES Purpose codes: W=taken from the wild, F=animals born in captivity, C=animals bred in captivity, R=ranched specimens.

| Rank | Species | No. of trophies | Mean (±SD) no. trophies/year | WOE | Mean (±SD) WOE/year | Exporter(s) | Source(s) |
| --- | --- | --- | --- | --- | --- | --- | --- |
| 1 | African elephant<br>*Loxodonta africana* | 538 | 24.45 ± 30.23 | 125 | 5.66 ± 5.08 | BW (166), ZW (162), ZA (114), TZ (56), MZ (23), NA (10), ZM (4), CM (3) | W (538) |
| 2 | American black bear<br>*Ursus americanus* | 485 | 22.04 ± 17.78 | 307 | 13.95 ± 8.32 | CA (474), US (11) | W (485) |
| 3 | Hippopotamus<br>*Hippopotamus amphibius* | 318 | 14.45 ± 16.56 | 91 | 4.12 ± 3.49 | TZ (96), ZM (95), ZW (61), UG (31), ZA (30), MW (2), MZ (2), CM (1) | W (318) |
| 4 | Hartman's mountain zebra<br>*Equus zebra hartmannae* | 260 | 11.81 ± 6.52 | 253 | 11.5 ± 6.48 | NA (219), ZA (41) | W (259), F (1) |
| 5 | Chacma baboon<br>*Papio ursinus* | 250 | 11.36 ± 9.62 | 244 | 11.09 ± 9.39 | ZA (165), NA (48), ZW (34) BW (2), ZM (1) | W (250) |
| 6 | Lion*<br>*Panthera leo* | 173 | 7.86 ± 5.97 | 162 | 7.35 ± 5.60 | ZA (100), TZ (28), ZW (17), NA (13), ZM (10), MZ (5) | W (89), C (83), F (1) |
| 7 | Caracal<br>*Caracal caracal* | 159 | 7.22 ± 5.39 | 159 | 7.22 ± 5.39 | ZA (153), NA (6) | W (158), F (1) |
| 8 | Southern lechwe<br>*Kobus leche* | 148 | 6.72 ± 4.76 | 148 | 6.72 ± 4.76 | ZA (81), ZM (36), BW (25), NA (6), | W (92), F (51), R (5) |
| 9 | Leopard<br>*Panthera pardus* | 122 | 5.45 ± 3.76 | 117 | 5.31 ± 3.49 | ZW (36), NA (23), TZ (21), MZ (15), ZM (15), BW (7), ZA (5) | W (122), C (1), I (1) |
| 10 | Nile crocodile<br>*Crocodylus niloticus* | 116 | 5.27 ± 5.34 | 102 | 4.63 ± 3.45 | ZA (43), ZW (32), ZM (19), TZ (11), MZ (9), NA (2) | W (112), C (4) |

\*Imports from the wild involved 4.04 ± 6.03 trophies/year (3.54 ± 5.5 WOE's/year), imports from animals bred in captivity involved 8.42 ± 2.43 trophies/year (8.42 ± 2.43 WOE's/year) and imports from animals born in captivity involved 0.04 ± .021 trophies/year (0.04 ± .021 WOE's/year).

Direct exports of hunting trophies from CITES-listed species from the UK in 2000-2021 involved 18 species, including 16 bird species and two (non-native) mammals (hog deer [*Axis porcinus*] and swamp deer [*Rucervus duvaucelii*]) (Supp. Material 2). Four bird species (pintail [*Anas acuta*], shoveler [*A. clypeata*], teal [*A. crecca*], and wigeon [*A. penelope*]), accounted for 98% of exports (945/968 trophies; 945 WOEs), which were sourced from the wild, and mainly imported by Malta (98%). This trade may not meet our definition of trophy hunting, but we included it because CITES purpose code H was used, i.e., the specimens were traded as hunting trophies. These species are no longer listed under CITES.

In 2015-2021, the UK imported an estimated 951 hunting trophies (738 WOEs) from 44 CITES-listed species and subspecies (Supp. Material 2). The top 10 species accounted for 78% of this trade, which mainly involved wild specimens, and included many of the same species for 2000-2021 (Table 2). Changes comprise the absence of leopard (*P. pardus*) and caracal (*Caracal caracal*) from this list but the inclusion of giraffe (*Giraffa camelopardalis*) and vervet monkey (*Chlorocebus pygerythrus*). Trade amounted to 135.85 ± 47.51 trophies/year (105.32 ± 40.58 WOEs/year). For the top 10 species this equates to between 5.57 ± 5.56 and 23.85 ± 16.67 trophies/year or 2.64 ± 1.27 and 15 ± 10.56 WOEs/year (Table 2). Lion imports mainly involved captive bred animals (83% or 59 trophies) and wild lion imports involved 1.71 ± 2.42 trophies/year (1.42 ± 1.71 WOEs/year). Of the 44 species and subspecies, 33 (75%), averaged <5 trophies imported annually, and 36 (82%) fewer than a mean of 5 WOEs a year. For 22 species and subspecies (50%) imports involved fewer than five trophies and for 25 species and subspecies (57%) less than one WOE a year.

**Table 2.** Top ten species imported to the UK as hunting trophies in 2015-2021 ranked by number of trophies, mean no. of trophies imported/year, WOEs imported, and mean WOEs imported/year. Source: CITES Trade Database (2023). WOEs=Whole organism equivalent. BW=Botswana, CA=Canada, MZ=Mozambique, NA=Namibia, TZ=Tanzania, ZA=South Africa, ZM=Zambia, ZW=Zimbabwe. CITES Purpose codes: W=taken from the wild, F=animals born in captivity, C=animals bred in captivity, R=ranched specimens.

| Rank | Species | No. of trophies | Mean (±SD) no. trophies/year | WOE | Mean (±SD) WOE/year | Exporter(s) | Source(s) |
| --- | --- | --- | --- | --- | --- | --- | --- |
| 1 | American black bear<br>*Ursus americanus* | 167 | 23.85 ± 16.67 | 71 | 10.14 ± 6.33 | CA (167) | W (167) |
| 2 | Chacma baboon<br>*Papio ursinus* | 105 | 15 ± 10.56 | 105 | 15 ± 10.56 | ZA (70), NA (20), ZW (14), ZM (1) | W (105) |
| 3 | African elephant<br>*Loxodonta africana* | 93 | 13.28 ± 12.28 | 34 | 4.82 ± 3.36 | ZW (60), ZA (15), BW (5), MZ (5), NA (4), TZ (2), ZM (2) | W (93) |
| 4 | Hartman's mountain zebra<br>*Equus zebra hartmannae* | 82 | 11.71 ± 6.47 | 75 | 10.71 ± 6.23 | NA (71), ZA (11) | W (82) |
| 5 | Lion*<br>*Panthera leo* | 71 | 10.14 ± 4.29 | 69 | 9.85 ± 3.71 | ZA (61), NA (6), MZ (2), TZ (2) | W (12), C (59) |
| 6 | Giraffe<br>*Giraffa camelopardalis* | 50 | 7.14 ± 11.17 | 25 | 3.57 ± 5.34 | ZA (43), ZW (5), NA (2) | W (50) |
| 7 | Southern lechwe<br>*Kobus leche* | 45 | 6.42 ± 5.41 | 45 | 6.42 ± 5.41 | ZA (34), NA (6), ZM (5) | F (32), W (11), R (2) |
| 8 | Vervet monkey<br>*Chlorocebus pygerythrus* | 45 | 6.42 ± 3.55 | 45 | 6.42 ± 3.55 | ZA (41), ZM (2), ZW (2) | W (43), R (2) |
| 9 | Nile crocodile<br>*Crocodylus niloticus* | 44 | 6.28 ± 4.75 | 43 | 6.14 ± 4.37 | ZA (33), ZW (5), MZ (2), NA (2), TZ (2) | W (43), C (1) |
| 10 | Hippopotamus<br>*Hippopotamus amphibius* | 39 | 5.57 ± 5.56 | 19 | 2.64 ± 1.27 | ZW (15), ZM (12), ZA (8), TZ (4) | W (39) |

\*Imports from the wild involved 1.71 ± 2.42 trophies/year (1.42 ± 1.71 WOE's/year) and imports from animals bred in captivity involved 8.42 ± 2.43 trophies/year (8.42 ± 2.43 WOE's/year).

Twenty-two countries exported hunting trophies to the UK in 2015-2021 but most exports were from four countries. South Africa, Canada, Namibia and Zimbabwe exported 94% of trophies in the top 10 species or 73% of all trophy imports to the UK. Seventy-nine percent of imports (753/951 trophies) were from countries where populations of the hunted species are stable, increasing, or abundant (Supp. Material 3). For example, the American black bear population in Canada is estimated at 450,000 individuals and is increasing (Garshelis *et al*. 2016). African elephant populations in Zimbabwe and South Africa number 82,000 and ∼27,000 animals, respectively, both increasing (Selier *et al*. 2016, CITES 2022). Hartmann’s Mountain zebra (*Equus zebra hartmannae*) numbers ∼44,000 individuals in Namibia and populations are increasing, as in South Africa (Gosling *et al*. 2019). Lion populations in South Africa have increased in recent decades and may be at carrying capacity (Bauer *et al*. 2015, 2016).

### 3.2 UK hunting trophy imports in context

For 2000-2021 the UK imported/exported 1929 CITES-listed animal species while species imported/exported as hunting trophies comprised <5% (91/1929 species) (Fig. 1). Combined imports to/exports from the UK in CITES-listed animal species in this period involved an estimated 4.93 million WOEs. The 3494 hunting trophies imported to the UK, involving an estimated 2549 WOEs, comprise <0.1% of UK trade in CITES-listed animal species. The same applies to 2015-2021; hunting trophies involved <4% of the species traded to/from the UK (44/1154 species) and <0.1% of UK trade in CITES-listed animal species. For comparison, many more species were traded, and trade volumes were greater, for commercial purposes, involving 1211 CITES-listed animal species and an estimated 4.85 million WOEs. Similarly, imports to/exports from the UK in animals as pets involved 568 species and an estimated 7752 WOEs, though most of these animals were reportedly from non-wild sources.

**Fig. 1.**
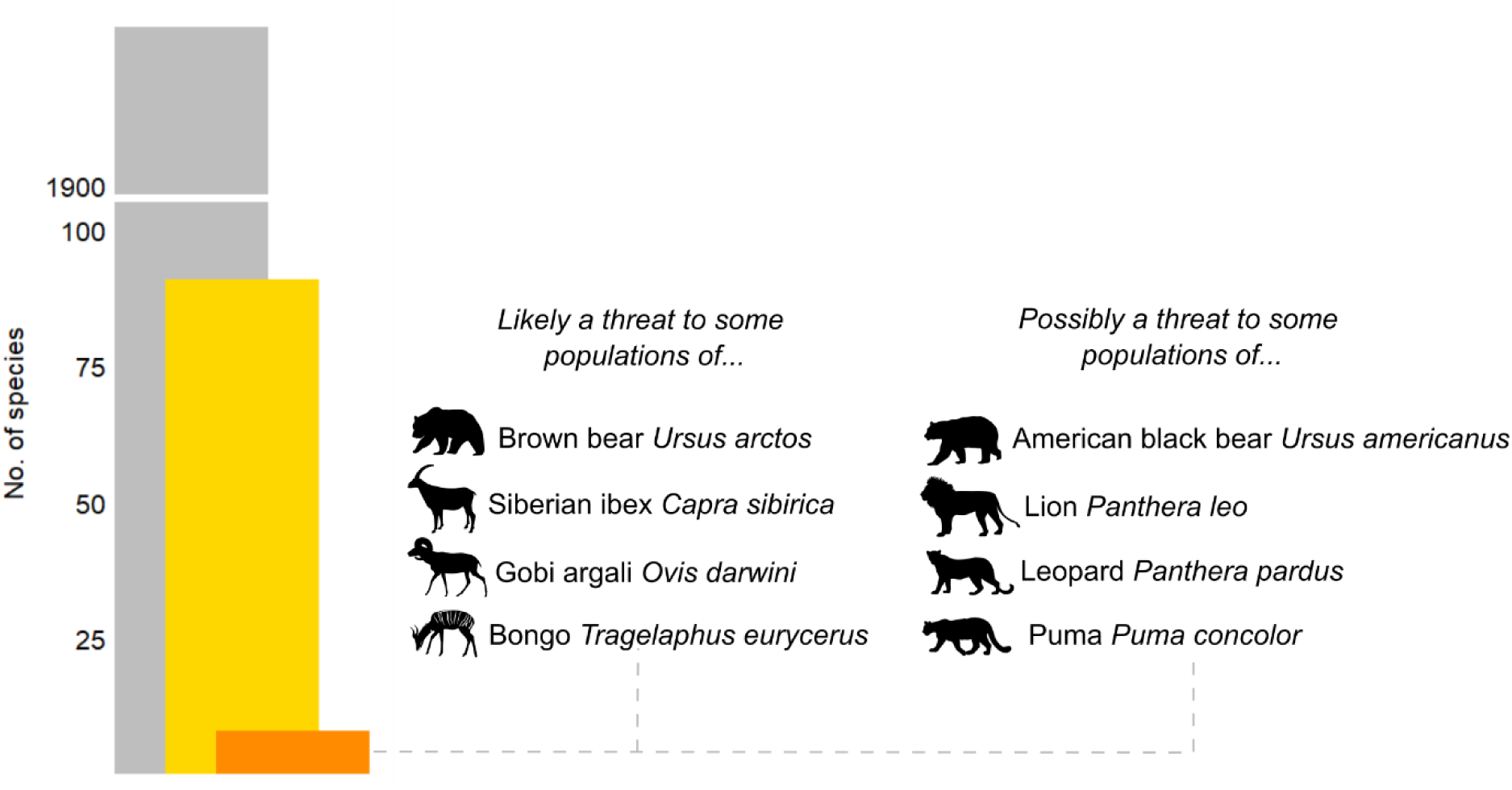
Number of CITES-listed animal species imported to/exported from the UK for all purposes (2000-2021) (grey), number of these species imported to/exported from the UK as hunting trophies in the same period (yellow), and number of these species for which trophy hunting is likely or possibly a local threat to some populations but does not contribute to the species being of elevated conservation concern (orange). Source: CITES Trade Database (2023) and IUCN Red List (2023).

### 3.3 Trophy hunting as a contributor to species being of elevated conservation concern and as a benefit provider

Of the species and subspecies imported to/exported from the UK as hunting trophies in 2000-2021, which were, or are, included under CITES, 60% (56/93 species) are categorised as Least Concern (LC) on the Red List, i.e., are not threatened with extinction (Supp. Material 4). We refer to 93 species because the Red List recognizes two distinct species of both urial and African elephant (Supp. Material 4). 40% (37 species) can be considered to be of elevated conservation concern (defined by IUCN as species assessed as Near Threatened, Threatened or Extinct in the Wild). Fourteen species (15%) are categorised as Near Threatened, 13 species (14%) as Vulnerable (VU), 5 (5%) as Endangered (EN), and 5 (5%) as Critically Endangered (CR) (Supp. Material 4). Intentional hunting and/or harvesting of aquatic resources is a major or minor threat to 77% of these taxa (Supp. Material 4), which consists of poaching and/or hunting that is illegal or poorly, or not, regulated.

Trophy hunting is not a major threat to any of the 93 species or subspecies imported/exported from the UK as hunting trophies in 2000-2021 (Supp. Material 4). However, it is likely a local threat (i.e., has or is causing localized declines) to some populations of 4% of these taxa (4 species) (Fig. 1). For brown bear (*Ursus arctos*) evidence suggests that where the species exists in large, contiguous populations, hunting rates are likely unsustainable in the short term but contribute to population fluctuations only rather than ongoing declines (McLellan *et al*. 2017). The Siberian ibex (*Capra sibirica*) is subject to regulated hunting for trophies in several countries, which targets males in the highest age classes (Reading *et al*. 2020a). If poorly regulated and managed, such hunting can have negative consequences, including on the sex and age composition of populations, but potential impacts are low if hunters only harvest a minor proportion of males (Michel and Rosen 2016, Reading *et al*. 2020a). For Gobi argali (*O. darwini*) evidence indicates that unsustainable trophy hunting comprises a minor localized threat in Mongolia (Reading *et al*. 2020b). The bongo (*Tragelaphus eurycerus*) is reportedly threatened by demand for hunting trophies, especially where hunting is poorly regulated (IUCN SSC Antelope Specialist Group 2016a), but this occurs in a small portion of the species’ range (Cameroon and Central African Republic) and represents, at most, a local threat.

Trophy hunting is possibly (rather than likely) a local threat to some populations of four species (4% of the 93 taxa) (Fig. 1). For lion, this may have at times contributed to population declines in Botswana, Namibia, Tanzania, Zimbabwe, Cameroon, and Zambia (Bauer *et al*. 2016). For leopard, where hunting for trophies over-concentrates on a particular area and targets animals in their prime that are reproductively active it can be detrimental to populations (Stein *et al*. 2020). Legal hunting of American black bear for trophies is well controlled in North America, but it may contribute to population fluctuations (Garshelis *et al*. 2016). The puma (*Puma concolor*) is legally hunted for trophies in many western and midwestern US states, which potentially comprises a minor, local threat (Neilsen *et al*. 2015).

For 20 species and subspecies imported to the UK as hunting trophies in 2000-2021, trophy hunting provides, or has the potential to provide, important benefits (Supp. Material 4). These include revenue generation for conservation, monetary and/or non-monetary benefits (e.g., meat and housing) to local communities, added value to wildlands that may be used for competing purposes such as agriculture, and enhanced population growth for threatened species. This includes species for which trophy hunting is likely or possibly a localized threat. Legal hunting for lion trophies has a net positive impact in some areas and is an important tool for conserving wild habitat and providing financial resources to governments and local communities (Bauer *et al*. 2016). Regulation which reduces the profitability of this hunting could result in widespread negative impacts for anti-poaching measures and the tolerance of lion outside protected areas (Hunter *et al*. 2013 cited in Bauer *et al*. 2016). Legal hunting of blesbok (*Damaliscus pygargus*) generates revenue and provides meat for local people (Dalton *et al*. 2019). For white and black rhino (*Diceros bicornis*), selective hunting of older males can increase population growth rates and provide resources for protection (Emslie 2020a, b). For bongo, well-regulated hunting for trophies has the potential to provide economic justification for preserving large areas of habitat in remote areas of Central Africa where possibilities for commercially successful tourism are limited (IUCN SSC Antelope Specialist Group 2016a). For Siberian ibex, banning legal hunting for trophies and/or associated trade risks removing incentives to prevent poaching, which may increase human-caused mortality of the species (Reading *et al*. 2020a).

### 3.4 Evaluating the UK Government’s impact assessment of the bill

We rated all five sections of the impact assessment weak because inappropriate assumptions were made, the quality of key analyses were poor, and/or there were areas that could be improved substantially to understand the likely impact of the bill. We discuss major areas of concern here, summarize key issues in Table 3 and the full evaluation is in Supp. Material 5.

**Table 3.** Summary of key issues and areas of improvement identified in the UK’s government’s impact assessment of the Hunting Trophies (Import Prohibition) Bill. Some issues apply to more than one section of the impact assessment. See Supp. Material 5 for the full evaluation.

| Section | Quality | Summary of key issues identified and areas of improvement |
| --- | --- | --- |
| 1. Policy rationale | Weak | The rationale focuses on importer-reported data only to characterize the UK’s role in the global trade in hunting trophies when exporter-reported quantities are often more complete, likely underestimating the UK’s role in this trade<br>The rationale assumes that the proposed legislation will disincentivize legal hunting of species included under the bill by people wishing to export hunting trophies to the UK, which may or may not be the case; empirical evidence referred to in Section 2 of the impact assessment suggests that this may not be the case and hunting may continue (though potentially at lower levels)<br>The assessment would be improved by robust analysis of the likely impacts of the bill on hunter behavior (e.g., would hunters plausibly continue to hunt, or not, and under what conditions)<br>The impact assessment focuses on the impacts of the proposed policy options in the UK, but the analysis could be improved by considering impacts in countries where trophy hunting takes place, especially as the impact assessment suggests that the highest costs would be expected to be borne by actors outside of the UK (e.g., local people and hunting operators)<br>The impact assessment could be improved by robust analysis of the threats to species that are subject to legal hunting for trophies for import into the UK, including explicit consideration of the magnitude of threat (e.g., major vs. minor), if any, and the scope, severity, and impact of any threat |
| 2. Costs and benefits | Weak | It is suggested that most costs can likely be expected to be borne by actors outside of the UK (e.g., hunting operators in countries where hunting trophies are acquired for import) but potential costs to such businesses are mentioned only briefly and are not quantified or monetised, though reference is made to the costs of some hunts<br>The environmental and societal impacts of the proposed policy options, in particular on countries and peoples beyond the UK, are considered vaguely. For example, “potential loss of income for communities that rely on trophy hunting” and “wildlife may lose part of its value due to no longer being economically competitive with other land uses, leading to negative conservation outcomes”<br>It is assumed that benefits will result from a reduction in the acquisition of hunting trophies and subsequent import to the UK as the species involved receive greater protection, which is misleading and not necessarily the case<br>The assumption that reducing trophy hunting will result in enhanced protection of target species does not hold and presumes that the institutional settings are appropriate and necessary resources available for protecting such species where hunting takes place but no justification is provided<br>For policy option #1 (Further restrict import and export of hunting trophies from Annex A and Annex B species), it is assumed that lost custom for hunting trips could be replaced by other forms of tourism, such as “eco-wildlife tourism” but this assumes that areas where species are legally hunted for trophies for import to the UK are suitable for these activities; prevailing evidence suggests that this is not the case for reasons including the remoteness of some areas where hunting takes place and lack of tourist infrastructure |
| 3. Risks and unintended consequences | Weak | The impact assessment would be improved by evidence that major exporting countries have been consulted on the proposed legislation, including insights on the potential impacts of the bill in those countries |
| 4. Wider impacts | Weak | It is suggested that a reduction in trophy hunting could result in positive knock-on effects for species numbers, biodiversity, and ecosystem services but this cannot be assumed because outcomes of the proposed policies could be positive or negative for species based on a range of social (e.g., the attitudes of local people), economic (e.g., incentives for legal and/or illegal offtake), and governance factors (e.g., the legitimacy of local laws and bans/restrictions imposed by importing countries) in areas where this hunting takes place<br>It is recognised in brief that banning the legal movement of hunting trophies could lead to increased illegal wildlife trade and reduce the amount of protein local communities receive as a by-product of trophy hunting but the impact assessment does not present detail on remedial measures beyond suggesting other forms of tourism (e.g., photographic tourism) and “other alternative wildlife management strategies” will be needed |
| 5. Post-implementation review | Weak | It is not clear whether proposed engagement with stakeholders during monitoring and evaluation includes individuals, groups, and businesses internationally (e.g., local communities and hunting operators in areas where trophy hunting takes place), which would be expected to bear most of the costs of the proposed legislation, or just those in the UK |

The impact assessment considered costs to UK individuals and businesses, but a major limitation is that it failed to adequately consider the costs and potential benefits to people (e.g., Indigenous people and local communities) in countries where trophy hunting takes place and helps sustain livelihoods (Table 3, Supp. Material 5). This is despite the implication in the assessment that most of the costs would likely be incurred by people who live in countries where trophy hunting takes place. The impact assessment would be markedly improved by explicit consideration of these costs, which would enable a more comprehensive characterization of the UK’s role in international hunting trophy trade and a better understanding of the likely impacts of the proposed policy.

The impact assessment assumed that the proposed legislation would disincentivize trophy hunting by people wanting to import trophies to the UK. This may or may not be the case but was weakly analyzed. There was little exposition of the actual impact that the proposed policy may have on hunter behavior and little supporting evidence was provided. The assessment would be much improved by robust analysis of the likely impacts of the bill on hunter behavior and whether it would plausibly disincentivize hunting, or not.

Additionally, the impact assessment assumed that the proposed policy would result in better protection of species with positive knock-on effects for biodiversity and ecosystem services. This may or may not be the case and cannot reasonably be assumed without a full analysis and supporting evidence. The policy could contribute to positive or negative outcomes for hunted species, related to social, economic, and/or governance factors in areas where hunting takes place (Table 3). Assuming species would be better protected presumes that the institutional arrangements are appropriate where hunting occurs and that the resources and revenue streams needed would be available, when they may not be, and evidence was not provided.

## 4. Discussion

The rationale for the Hunting Trophies (Import Prohibition) Bill was to ensure that imports of hunting trophies to the UK are not placing additional pressure on species of conservation concern, by prohibiting such imports from a reported ∼7000 species (UK Government 2021a). Our analyses indicate that the UK plays a minor role in the global trade of hunting trophies of CITES-listed species, and that hunting trophies comprise <0.1% of UK trade in all CITES-listed animal species. Although the UK imported hunting trophies from 73 CITES-listed species and subspecies in 2000-2021, these represented only 1% of the species included under the bill. Based on the Red List, trophy hunting is not a major threat contributing to any of these species being of elevated conservation concern. In 2015-2021, 79% of imports were from countries where populations of the hunted species are stable increasing, or abundant, including species for which trophy hunting is a likely or possible local threat. Imports of American black bear, brown bear, Siberian ibex, and lion trophies were from countries with healthy populations. Lion imports involved <2 wild animals a year, mainly from countries for which evidence suggests populations are increasing. The leopard is perhaps an exception because although imports were low (<4 animals/year), all were from sub-Saharan Africa where many populations are declining, or their status is unknown (though quotas are typically set by government agencies using more detailed local information and adaptive management). Assuming past trade is indicative of future imports, the argument that the bill would reduce pressure on many threatened species subject to legal hunting for trophies is unfounded. Other threats, notably poaching and/or retaliatory killing, are much greater for most species imported to the UK as hunting trophies. Trophy hunting is likely or possibly a local threat to population of nine species, but there are likely more effective mitigation measures than the UK banning hunting trophy imports. For example, improved management at national and sub-national levels.

The bill could also undermine conservation efforts that are supported by trophy hunting. More evidence is needed on the potential impacts of the bill, but it could reduce revenue for conservation programs which rely on such hunting to fund wildlife management. Reduced funding could jeopardize law enforcement, anti-poaching efforts, and monitoring thereby increasing other threats to species and habitats. The bill could have negative, even devastating, impacts on Indigenous people and local communities who rely on such hunting for monetary and/or non-monetary benefits (e.g., meat and employment) (IUCN 2016a, Angula *et al*. 2018, Parker *et al*. 2023). Benefits to local communities vary (e.g., 0-100% of revenue generated) but are frequently very important to them (IUCN 2016a). For example, in Namibia conservation hunting generated USD 1.3 million and provided 326,000 kg of game meat in 2021, which were distributed to local communities (MEFT/NACSO 2022). Ultimately, the bill could influence the viability of conservation areas to conserve biodiversity with wildlife habitat being lost to competing land uses such as agriculture and mining (Lindsey *et al*. 2007, IUCN 2016a, Strampelli *et al*. 2022). In this context, the bill may contravene principles in the UK Environment Act (2021), specifically that UK policymaking should prevent environmental harm because the bill could contribute to more harm than it prevents. There is also a risk that the UK sets a precedent that other countries follow resulting in even greater harm to wildlife conservation efforts.

The rationale for the bill is also to respond to the UK Government’s hunting trophies consultation. 84% of respondents to the consultation indicated a preference to prohibit all hunting trophies entering or leaving the UK; however, 68% of all responses were linked to advocacy group campaigns (UK Government 2021b). Another poll found that fewer than half of Britons wanted a ban if it would harm conservation or local communities (Survation 2021). Recent research suggests that UK public opinion is more supportive of hunting programs that provide tangible benefits to people who live in hunting areas (e.g., meat and economic development; Hare *et al*. 2024). Many respondents to the consultation appear strongly opposed to trophy hunting likely because they consider it ethically unacceptable. This does not mean that prohibiting imports of hunting trophies is the most appropriate policy. In democratic societies public opinion should be considered in public policymaking but, critically, it should not be decisive but interpreted with other evidence using appropriate analytical capacity (Howlett *et al*. 2020). A key challenge for public policymaking is formulating proportionate policies based on all relevant bodies of evidence, including public opinion shaped by moral values. Regarding this bill, our analyses indicate that it is disproportionate and would be unlikely to achieve its intended effects while risking negative impacts on wildlife and local communities.

## 5. Recommendations

Recognizing that trophy hunting can benefit species but can have negative impacts if poorly regulated (Hare *et al*. 2023, IUCN 2012), what are alternative policy options to the proposed legislation? Several options exist (Supp. Material 6), which we argue are more proportionate and targeted than an indiscriminate ban on hunting trophy imports to the UK:

1. Do nothing differently. The UK could continue to implement rigorously the WTRs and EUWTRs (Northern Ireland), ensuring that imports and exports of hunting trophies of all Annex A and six Annex B species are based on robust NDFs and legal acquisition findings.
2. Remove the personal and household effects derogation from all Annex B species traded as hunting trophies. This would require import permits for all hunting trophies imported to the UK using CITES purpose code H enabling further assessment of sustainability, for example, based on requirements in CITES Resolutions on hunting trophies (Res. Conf. 17.9) and NDFs (Res. Conf. 16.7 [Rev. CoP17]).
3. Apply stricter measures to the import of hunting trophies from particular species; such measures are used for rhinos, bears (Ursidae spp.) and tiger (*P. tigris*) (UK Government 2023).
4. Implement a smart ban (Webster *et al*. 2022), analogous to the proposed conservation amendment to the Hunting Trophies (Import Prohibition) Bill (Fleming 2024). This would prohibit the import of hunting trophies except in circumstances where the benefits of this hunting tangibly contribute to the conservation of the hunted species and their habitat, there is an equitable sharing of hunting revenues with local communities, an adaptive management and monitoring system is in place, and the hunting area has good governance (Fleming 2024, see IUCN 2012).

If UK policymakers are committed to legislating on hunting trophy imports, we argue that a smart ban would be the most appropriate and evidence-based policy. It would raise the standard and scope of regulation without unduly affecting hunting for trophies where it is well-regulated and benefits species and Indigenous people and local communities. Under such a smart ban an import permit would be required for hunting trophies from species in Annex B of the WTRs, and imports of trophies from all species on Annexes A and B of the WTRs would be required to demonstrate conservation and other benefits as outlined, which would be legally binding, to qualify for an import permit (Fleming 2024). These measures would be stricter than those under CITES, and we would encourage the UK government to consult key exporting countries prior to enacting any law as recommended in CITES Res. Conf. 6.9 (Rev. CoP17) and Res. Conf. 17.9. Research suggests this policy may also reflect public opinion in the UK more accurately than an indiscriminate ban (Hare *et al*. 2024).

Public policy to address biodiversity loss requires context-specific solutions (Ostrom 2007, IPBES 2022). Our analyses suggest that an indiscriminate ban on imports of hunting trophies to the UK would be disproportionate and may harm biodiversity and rural livelihoods, in part because it does not differentiate between different types of legal hunting across social-ecological and governance contexts (Hare *et al*. 2023). Crucially, the UK Government’s impact assessment failed to adequately consider the likely impacts of this policy on people outside of the UK who are expected to incur most of the costs. The UK Government may consider it disproportionate to evaluate such costs, but we argue that if policymakers were serious about conserving biodiversity they would refrain from proposing oversimplistic policy solutions to complex biodiversity issues. We recommend that UK Government impact assessments concerning biodiversity internationally go beyond UK people and businesses and consider international impacts. This should involve consultation with relevant countries to examine the costs and benefits of policy options, which would help ensure that future policy is appropriately evidence-based (Sutherland 2020), and has the greatest likelihood of benefitting biodiversity, while avoiding environmental and socio-economic harm.

## Supplementary Material

### Supplementary Material 1 – Methods

#### Analyzing international trade in CITES-listed species

##### Number of hunting trophies from CITES-listed species and WOEs traded globally

We downloaded a comparative tabulation report for 2000-2021 including all exporting and importing countries, all sources, for the purpose of “hunting trophy” (purpose code H), and all trade terms and taxa. We analyzed direct trade only because it comprised the vast majority of the trade by volume. In estimating the number of hunting trophies and WOEs traded we excluded trade using the term “live”. We subsequently ranked importers and exporters based on reported trade volumes using the number of trophies. We then calculated the whole organism equivalents (WOEs) for this trade adapting the approach by Harfoot *et al*. (2018). This entailed converting products in trade to WOEs, for example 5 bodies represent 5 WOEs and 4 feet represent 1 mammal WOE (Table S1). To estimate WOEs we used the terms in Table S1 with unit =blank, number of specimens, belly skins, and hornback skins. To estimate the number of trophies and WOEs traded in 2015-2021, we summarized data for this period. As the CITES trade data may refer to permits issued rather than number of specimens traded, and as estimating the number of WOEs may count animal parts from the same animal more than once, our estimates of trade volumes and WOEs may be overestimates.

##### The UK’s role in the international trade of hunting trophies in CITES-listed species

We downloaded a comparative tabulation report for the period 2000-2021 for all exporting countries with the UK as the importing country, including all sources, for the purpose of “hunting trophy” (purpose code H), and all trade terms and all taxa. To estimate the number of hunting trophies in trade we used all trade terms (except “live”) and units =blank and number of specimens but not other units (gm and kg). We visually inspected the data and added small volumes of trade reported by importers to our summaries where such trade was evidently not reported by exporters being careful to avoid double counting (CITES Secretariat and UNEP-WCMC 2022). As above, we estimated the number of WOEs in trade as hunting trophies adapting the approach by Harfoot *et al*. (2018). To estimate WOEs we used only those terms included in Table S1 with unit=blank and number of specimens. We repeated this process for exports of hunting trophies from the UK, downloading data using the same terms as above but with the UK as the exporting country and all other countries as importers. We did so recognizing that the UK may export hunting trophies as well as being an importer. We also repeated the process described above to estimate WOEs for UK exports. To estimate the number of trophies and WOEs traded in 2015-2021, we summarized data for this period. The caveats relating to CITES trade data highlighted above also apply here.

**Table S1.** Conversion factors used to estimate WOEs for hunting trophies. Adapted from Harfoot *et al*. (2018).

| Class | Term | Description | Factor to equate to WOE |
| --- | --- | --- | --- |
| Aves | BOD | Bodies | 1 |
|  | SKE | Skeleton | 1 |
|  | SKI | Skins | 1 |
|  | SKU | Skulls | 1 |
|  | TAI | Tails | 1 |
|  | TRO | Trophies | 1 |
| Mammalia | BOD | Bodies | 1 |
|  | FOO | Feet | 0.25 |
|  | HOR | Horns | 1 |
|  | SKE | Skeleton | 1 |
|  | SKI | Skins | 1 |
|  | SKU | Skulls | 1 |
|  | TAI | Tail | 1 |
|  | TEE | Teeth – *Hippopotamus amphibius* only | 0.083 |
|  | TRO | Trophies | 1 |
|  | TUS | Tusks – *Loxodonta africana* only | 0.532 |
| Reptilia | BOD | Bodies | 1 |
|  | CAR | Carapace | 1 |
|  | FOO | Feet | 0.25 |
|  | SKE | Skeleton | 1 |
|  | SKI | Skins | 1 |
|  | SKU | Skulls | 1 |
|  | TRO | Trophies | 1 |

##### UK trade in CITES-listed animal species overall, for commercial purposes, and as pets

To contextualize the number of CITES-listed species imported to/exported from the UK as hunting trophies among CITES-listed animal species traded overall and for commercial purposes and as pets, including associated trade volumes, we downloaded two comparative tabulation reports. The first comprised data on all exporting countries with the UK as the importing country, including all sources, purposes, trade terms, and taxa for the period 2000-2021. The second included the same terms but with the UK as the exporter and all other countries as importers. We calculated the number of species imported to/exported from the UK for the periods 2000-2021 and 2015-2021. To estimate trade volumes we calculated the number of WOEs imported to and exported from the UK overall and then for both commercial purposes (purpose code T) and as pets (purpose codes P and B; see below) using the terms in Table S2 and units=blank and number of specimens and totaled imports and exports. To estimate WOEs for trade in pets, we used purpose codes P (personal) and B (breeding in captivity), the terms live, eggs, and eggs-live, and units=blank and number of specimens. The caveats relating to CITES trade data highlighted above also apply here.

**Table S2.**
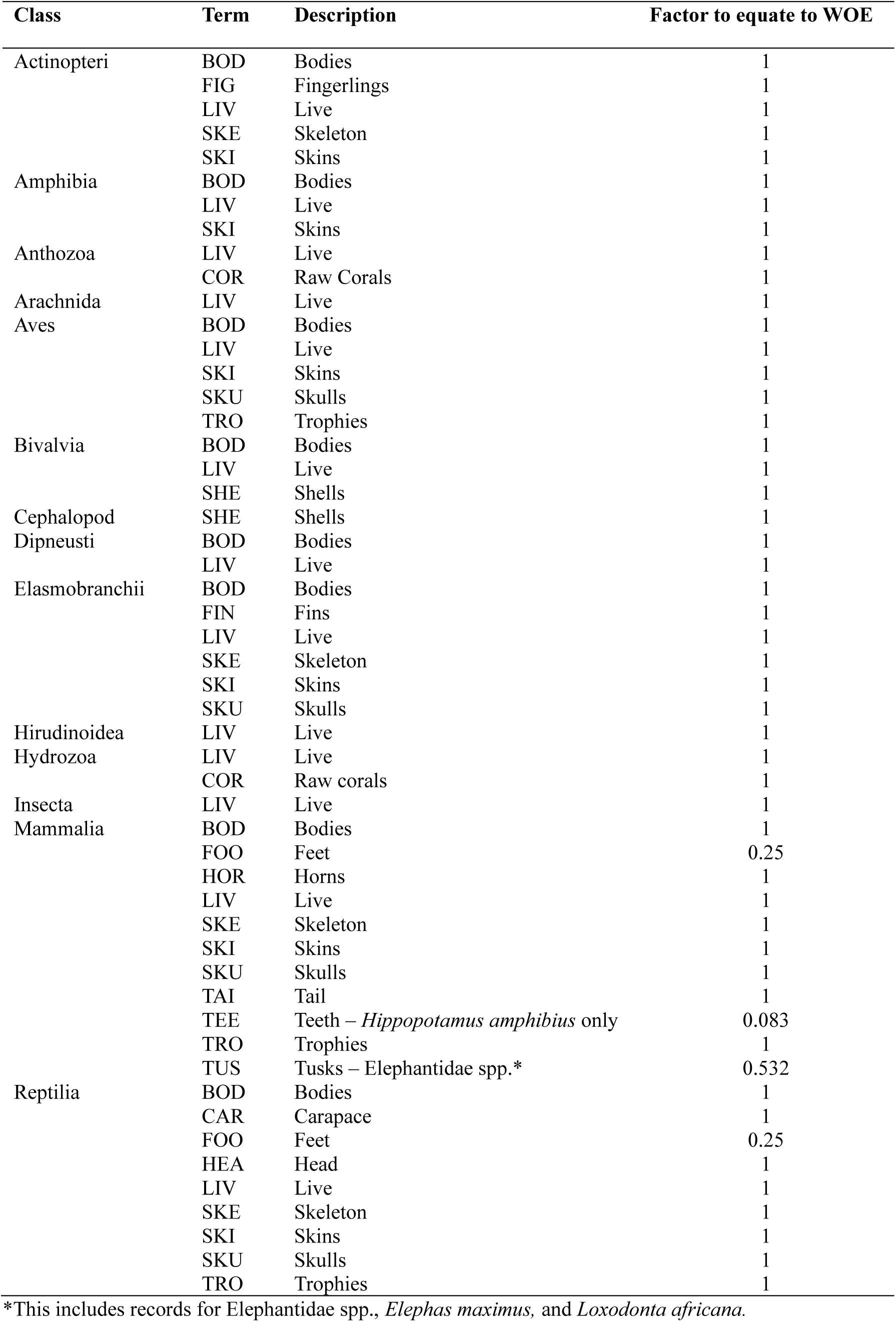
Conversion factors used to estimate WOEs adapted from Harfoot *et al*. (2018).

#### Trophy hunting as a contributor to species being of elevated conservation concern and as a benefit provider

##### Species database

Our MS Excel database includes data from IUCN Red List assessments (using Red List version 2022-2) and Species+ (UNEP 2023) for species directly imported to/exported from the UK as hunting trophies in the period 2000-2021^1^. The database contains the following fields: class, common name, scientific name, CITES Appendix, date of last assessment, Red List Category, population size, population trend, and whether intentional hunting of the species is a current threat (yes/no) based on the following threat codes: 5.1.1. (Hunting & collecting terrestrial animals → Intentional use (species being assessed is the target)), 5.4.1. (Fishing & harvesting aquatic resources → Intentional use: subsistence/small scale (species being assessed is the target)), and/or 5.4.2. (Fishing & harvesting aquatic resources → Intentional use: large scale (species being assessed is the target)). We included a field for which of these threat codes had been applied to species (e.g., 5.1.1.) and the timing of those threats (e.g., past or ongoing). For more information on how the Red List generally defines major threats see IUCN (2023). We included a field that contained the narrative text from the threats field and justification field from Red List assessments. We also included a field containing relevant information from the Use and Trade section of each assessment (where information was available). The Use and Trade Classification Scheme includes a specific category (“Sport hunting/specimen collecting”) that provided a helpful countercheck, noting that there is no direct relation between the use and trade scheme and the threat scheme (i.e., the Use and Trade classification scheme is intended to document use at any level whether a threat or not). The classification of species was undertaken by one author and the results were discussed with and/or checked by three other authors. The Red List was updated in December 2023, and we checked and updated information in our database against Red List version 2023-1 (IUCN 2024).

Recognizing that trophy hunting can provide benefits to species and local people, we also recorded in our database whether Red List assessments for species referred to such benefits, either actual or potential. To do so we also read the Conservation Actions section of assessments and considered species to benefit (actually or potentially) where this was stated in the selected sections of assessments reviewed.

##### Species examples

The giraffe (*Giraffa camelopardalis*) was categorised as not having trophy hunting as a major threat contributing to the species being listed as Vulnerable. This is because, while intentional use is a major threat, there is no evidence that this is driven by legal hunting for trophies.

Indeed, the Red List account makes it clear that the threat is illegal hunting. The Gobi argali (*Ovis darwini*) was categorised as trophy hunting *likely* comprising a local threat (but does not contribute to elevated conservation concern) because this is stated in the threats narrative in the assessment for this taxon. The lion (*Panthera leo*) was categorised as trophy hunting *possibly* comprising a local threat. This is because the narrative text in the assessment states that this hunting “*may have at times*” contributed to population declines.

Regarding benefits, the black rhino (*Diceros bicornis*) was considered to benefit from trophy hunting because the Conservation Actions section of the assessment states that “*sport hunting quotas have been approved for the two Range States with biggest populations (South Africa and Namibia). Removal of specific individuals can enhance demographic and/or genetic conservation*.” The Siberian ibex (*Capra sibirica*) was considered to benefit from trophy hunting because hunters purchase hunting licenses from the government and an identified conservation action is to “*begin using the money generated from trophy hunting to pay for conservation and management of the species, ideally using community-based approaches*.”

##### Limitations to using IUCN Red List data

Further to the main text, we acknowledge that our assessment of the degree to which trophy hunting is a major threat or not is based on a direct interpretation of information contained in the Red List which may not be current or complete. As such we cannot, and do not, purport to definitively state that trophy hunting is not a major threat to any species; instead we note that, based on the best available data in the Red List, this does not currently appear to be the case. We also note that for the 93 species in our dataset only one Red List account (Collared peccary *Pecari tajacu*) is outdated (i.e., older than 10 years) though 12 assessments were conducted in 2014 and 7 in 2015.

#### Evaluating the UK Government’s impact assessment of the bill

Regulatory proposals in the UK are accompanied by impact assessments, which evaluate the likely associated risks, costs, and benefits of proposed regulation, and impacts on businesses, charities and voluntary organisations, the public sector, and individuals (Regulatory Policy Committee, n.d.). Assessments are based on the ROAMEF policy cycle (Rationale, Objectives, Appraisal, Monitoring, Evaluation, and Feedback) and use cost-benefit analyses (Department for Business, Energy and Industrial Strategy, 2020). Guidance is provided to government departments on conducting impact assessments through the Better Regulation Framework (Department for Business, Energy and Industrial Strategy, 2020) and HM Treasury’s Green Book - Central Government Guidance on Appraisal and Evaluation (hereafter “Green Book”; UK Government 2022).

Impact assessments should clearly explain the issue the government is trying to tackle, how they intend to tackle it, and what the effects of proposals will be (Department for Business, Energy and Industrial Strategy, 2020). The Better Regulation Framework and Green Book detail how to appraise potential policy options, including associated justification, and how to make the case for preferred policy options. Key considerations include that policy development should be based on objective evidence; assumptions, where needed, are reasonable and justified by transparent reference to the research on which they are based; that risks associated with the regulation are understood; and, that policy development should consider the impact of proposed regulation on specific groups in society, places, or businesses, including unintended consequences. Where effects of regulation are significant for particular groups, analysis identifying gaining and losing groups and estimates of the effects should be undertaken. The Green Book also states that impacts on natural capital should be considered, including explicit consideration of wildlife and the provision of ecosystem services. The principal focus of risk assessments is impacts in the UK but the Green Book states that it is sometimes reasonable to consider the costs and benefits to people living outside the UK, and where appraisal of Official Development Assistance (ODA) is concerned, this should include the costs and benefits to the recipient countries.

Based on the estimated equivalent annual net direct cost to business (EANDCB) different types of impact assessment are required (Department for Business, Energy and Industrial Strategy, 2020). If the proposed regulation involves EANDCB of £±5 million, full impact assessments are required, adhering to the aforementioned guidance, and scrutiny by the Regulatory Policy Committee (RPC) is required. In circumstances where the EANDCB of proposed regulation is <£5 million, only a proportionate analysis is required (Department for Business, Energy and Industrial Strategy, 2020). The RPC is an independent regulatory body for the UK Government, which assesses the quality of evidence and analysis used to inform qualifying government regulatory proposals based on the aforementioned guidance but does not provide opinions on policies themselves (Regulatory Policy Committee, n.d.). On evaluating impact assessments, the RPC formally rates them as green (=fit for purpose) or red (=not fit for purpose). Fit for purpose impact assessments are those where there are no significant concerns over the quality of the assessment or there are minor issues that could be improved. Impact assessments that are not fit for purpose are those where there are major concerns over the quality of the evidence and analysis and the overall quality of the assessment. The RPC recently introduced quality indicators, ranging from *very weak* to *good*, based on a range of criteria, and covering key areas of impact assessments (e.g., “rationale and options”) but which are not formally rated (Regulatory Policy Committee 2021).

For proportionate analysis of regulatory impact assessments, guidance indicates that the level of analysis should be proportionate to the problem the regulation is addressing and reflect the scale or impact of the measure(s) (Department for Business, Energy and Industrial Strategy, 2020). This includes proportionate cost-benefit analysis to inform decision-making. For proportional analyses, scrutiny of impact assessments by the RPC is optional.

Using the RPC’s quality indicators (Table S3), we evaluated the UK Government’s regulatory impact assessment of the Hunting Trophies (Import Prohibition) Bill. The impact assessment for this bill did not qualify for, and was not, scrutinized by the RPC. Guidance on conducting impact assessments states that assessments should focus on the UK but may at times focus on costs and benefits outside of the country. As such we evaluated the impact assessment considering costs and benefits both within and outside of the UK, recognizing the international nature of trade in hunting trophies of CITES-listed species. This decision was informed by an initial read of the impact assessment prior to analysis, which indicated that such impacts had, at least, been acknowledged. We evaluated each section of the impact assessment by reading it carefully and noting both positive and negative aspects with reference to the criteria (Table S3). We did not provide an overall rating for the assessment.

Finally, recognizing calls for researchers to reflect on their position in research (Blair 2016), we recognize that as an author group, we have engaged in research and policy formulation relating to wildlife conservation and management, including research and monitoring of legal hunting of wildlife for trophies specifically, for periods of up to several decades each and therefore have familiarity with the subject area.

**Table S3.**
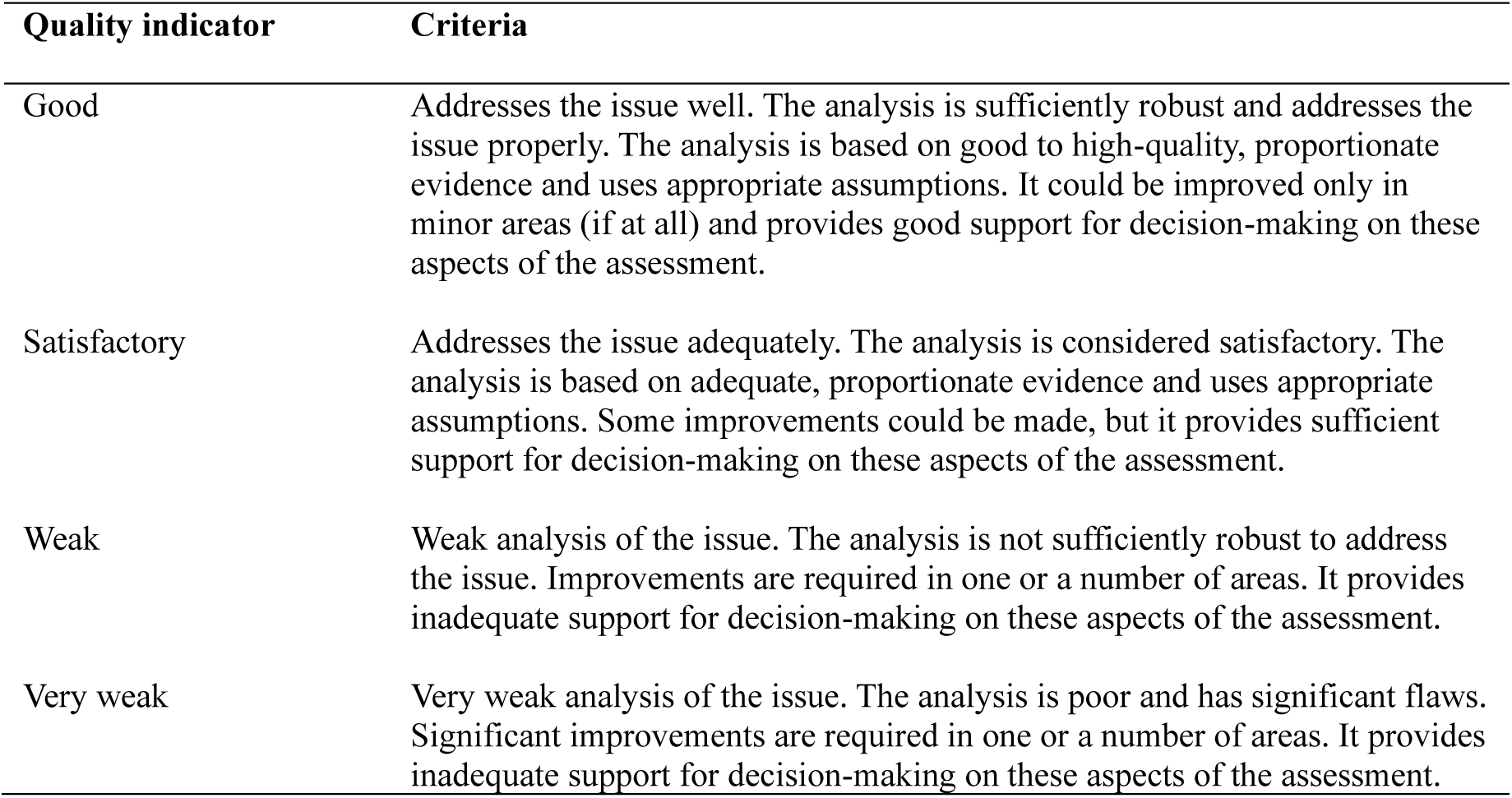
Quality indicators used to evaluate the impact assessment for the Hunting Trophies (Import Prohibition) Bill. Source: Regulatory Policy Committee (2021).

### Supplementary Material 2 – Results – International trade in hunting trophies

**Table S4.** Top 20 exporters of hunting trophies from CITES-listed species (2000-2021) based on number of trophies.

| Rank | Exporter | Estimated number of hunting trophies |
| --- | --- | --- |
| 1 | Canada | 140,954 |
| 2 | South Africa | 132,341 |
| 3 | Mozambique | 46,032 |
| 4 | Namibia | 45,715 |
| 5 | Zimbabwe | 35,321 |
| 6 | Bulgaria | 33,155 |
| 7 | Zambia | 29,109 |
| 8 | Tanzania | 21,092 |
| 9 | Botswana | 19,190 |
| 10 | Russian Federation | 13,696 |
| 11 | Mexico | 10,789 |
| 12 | Argentina | 7921 |
| 13 | United States | 3972 |
| 14 | Cameroon | 2795 |
| 15 | Kyrgyzstan | 2595 |
| 16 | Romania | 2330 |
| 17 | Central African Republic | 1737 |
| 18 | Mongolia | 1550 |
| 19 | United Kingdom | 968 |
| 20 | Pakistan | 801 |

**Table S5.**
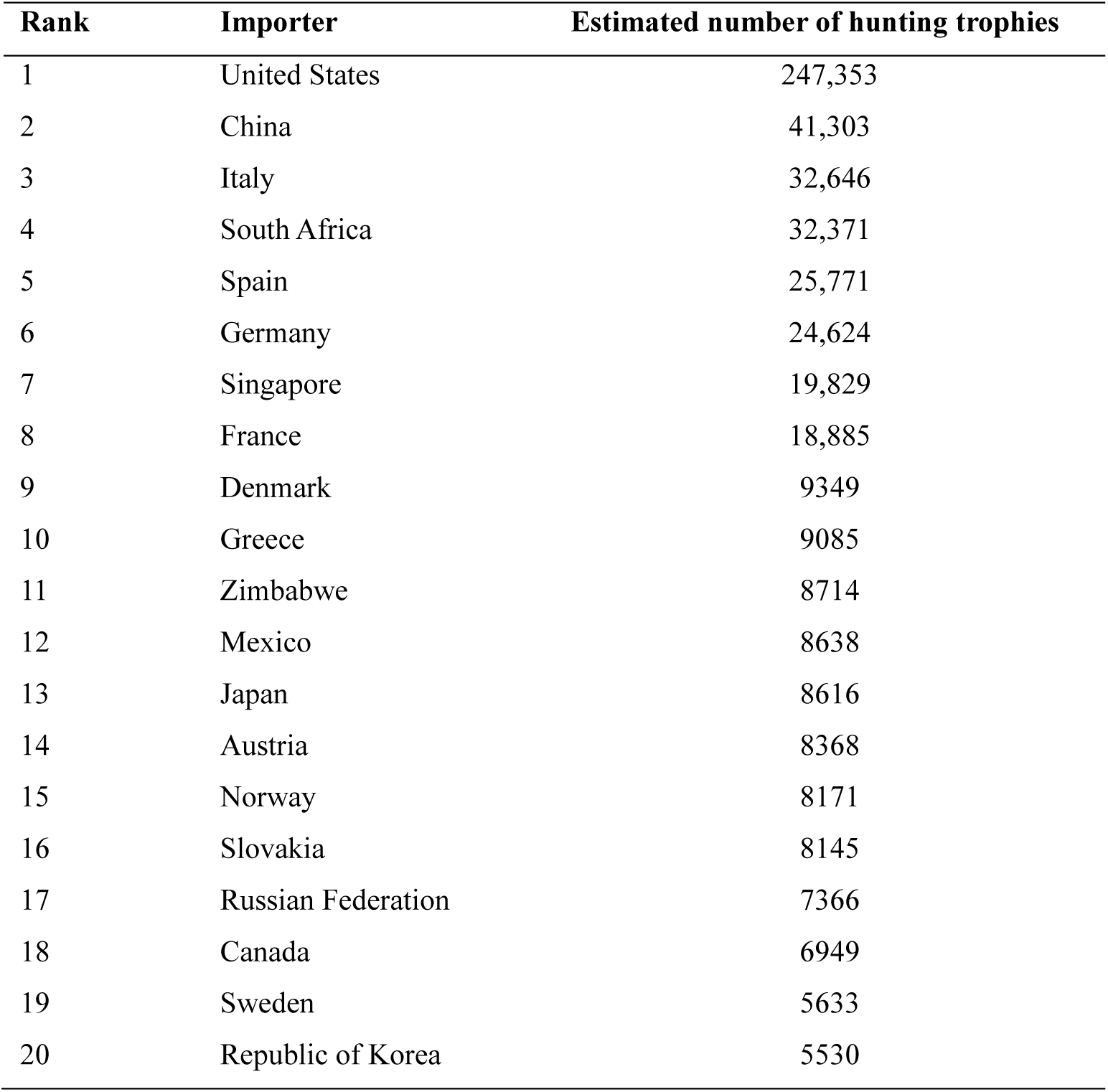
Top 20 importers of hunting trophies from CITES-listed species (2000-2021) based on number of trophies.

**Table S6.**
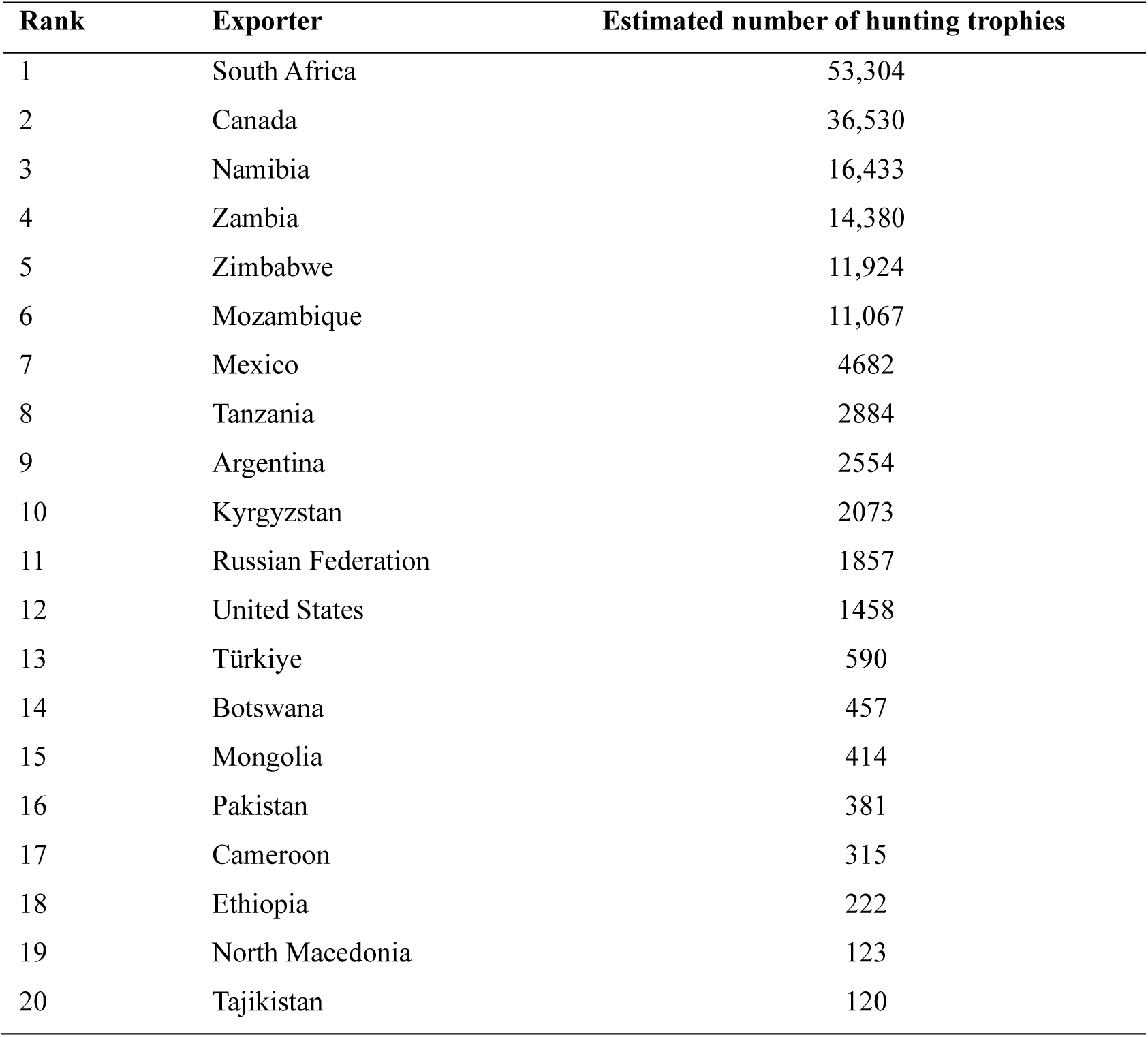
Top 20 exporters of hunting trophies from CITES-listed species (2015-2021) based on number of trophies.

| Rank | Exporter | Estimated number of hunting trophies |
| --- | --- | --- |
| 1 | South Africa | 53,304 |
| 2 | Canada | 36,530 |
| 3 | Namibia | 16,433 |
| 4 | Zambia | 14,380 |
| 5 | Zimbabwe | 11,924 |
| 6 | Mozambique | 11,067 |
| 7 | Mexico | 4682 |
| 8 | Tanzania | 2884 |
| 9 | Argentina | 2554 |
| 10 | Kyrgyzstan | 2073 |
| 11 | Russian Federation | 1857 |
| 12 | United States | 1458 |
| 13 | Türkiye | 590 |
| 14 | Botswana | 457 |
| 15 | Mongolia | 414 |
| 16 | Pakistan | 381 |
| 17 | Cameroon | 315 |
| 18 | Ethiopia | 222 |
| 19 | North Macedonia | 123 |
| 20 | Tajikistan | 120 |

**Table S7.** Top 20 importers of hunting trophies from CITES-listed species (2015-2021) based on number of trophies.

| Rank | Importer | Estimated number of hunting trophies |
| --- | --- | --- |
| 1 | United States | 74,107 |
| 2 | China | 39,791 |
| 3 | South Africa | 10,524 |
| 4 | Slovakia | 6794 |
| 5 | Germany | 6501 |
| 6 | Kyrgyzstan | 5391 |
| 7 | Singapore | 5188 |
| 8 | Japan | 4632 |
| 9 | Spain | 4092 |
| 10 | France | 3111 |
| 11 | Denmark | 2991 |
| 12 | Canada | 2987 |
| 13 | Russian Federation | 2943 |
| 14 | Italy | 2737 |
| 15 | Sweden | 2464 |
| 16 | Austria | 2421 |
| 17 | Mexico | 2316 |
| 18 | Norway | 2066 |
| 19 | Hungary | 1922 |
| 20 | Zimbabwe | 1764 |

**Table S8.** Species and subspecies (n=73) imported to the UK as hunting trophies (2000-2021), estimated number of trophies, mean (± SD) number of trophies traded/year, WOEs traded in this period, and mean (± SD) WOEs traded in this period/year, ranked by number of trophies.

| Rank | Common name | Scientific name | Estimated trophies | Mean ± SD | WOE | WOE (mean ± SD) |
| --- | --- | --- | --- | --- | --- | --- |
| 1 | African elephant | *Loxodonta africana* | 538 | 24.45 ± 30.23 | 125 | 5.66 ± 5.08 |
| 2 | American black bear | *Ursus americanus* | 485 | 22.04 ± 17.78 | 307 | 13.95 ± 8.32 |
| 3 | Hippopotamus | *Hippopotamus amphibius* | 318 | 14.45 ± 16.56 | 91 | 4.12 ± 3.49 |
| 4 | Hartmann's mountain zebra | *Equus zebra hartmannae* | 260 | 11.81 ± 6.52 | 253 | 11.5 ± 6.48 |
| 5 | Chacma baboon | *Papio ursinus* | 250 | 11.36 ± 9.62 | 244 | 11.09 ± 9.39 |
| 6 | Lion* | *Panthera leo* | 173 | 7.86 ± 5.97 | 162 | 7.35 ± 5.60 |
| 7 | Caracal | *Caracal caracal* | 159 | 7.22 ± 5.39 | 159 | 7.22 ± 5.39 |
| 8 | Southern lechwe | *Kobus leche* | 148 | 6.72 ± 4.76 | 148 | 6.72 ± 4.76 |
| 9 | Leopard | *Panthera pardus* | 122 | 5.45 ± 3.76 | 117 | 5.31 ± 3.49 |
| 10 | Nile crocodile | *Crocodylus niloticus* | 116 | 5.27 ± 5.34 | 102 | 4.63 ± 3.45 |
| 11 | Grey wolf | *Canis lupus* | 100 | 4.54 ± 9.74 | 52 | 2.36 ± 2.17 |
| 12 | Brown bear | *Ursus arctos* | 99 | 4.5 ± 3.71 | 99 | 4.5 ± 3.71 |
| 13 | Vervet monkey | *Chlorocebus pygerythrus* | 79 | 3.59 ± 3.94 | 79 | 3.59 ± 3.94 |
| 14 | Hamadryas baboon | *Papio hamadryas* | 75 | 3.4 ± 4.99 | 74 | 3.36 ± 4.94 |
| 15 | Giraffe | *Giraffa camelopardalis* | 50 | 2.27 ± 6.87 | 25 | 1.13 ± 3.32 |
| 16 | Topi | *Damaliscus lunatus* | 43 | 1.95 ± 3.52 | 43 | 1.95 ± 3.52 |
| 17 | Grivet monkey | *Chlorocebus aethiops* | 42 | 1.9 ± 3.08 | 42 | 1.9 ± 3.08 |
| 18 | Bontebok | *Damaliscus pygargus pygargus* | 41 | 1.86 ± 2.18 | 41 | 1.86 ± 2.18 |
| 19 | Afro-Asiatic wildcat | *Felis silvestris* | 38 | 1.72 ± 2.76 | 38 | 1.72 ± 2.76 |
| 20 | Blackbuck | *Antelope cervicapra* | 28 | 1.27 ± 1.57 | 28 | 1.27 ± 1.57 |
| 21 | Yellow baboon | *Papio cynocephalus* | 25 | 1.13 ± 1.95 | 25 | 1.13 ± 1.95 |
| 22 | Southern white rhino | *Ceratotherium simum simum* | 22 | 1 ± 1.66 | 16 | 0.72 ± 1.12 |
| 23 | African civet | *Civettictis civetta* | 22 | 1 ± 1.15 | 22 | 1 ± 1.15 |
| 24 | Puma | *Puma concolor* | 21 | 0.95 ± 1.29 | 21 | 0.95 ± 1.29 |
| 25 | Serval | *Leptailurus serval* | 20 | 0.9 ± 1.06 | 20 | 0.9 ± 1.06 |
| 26 | Siberian ibex | *Capra sibirica* | 19 | $0.86 \pm 2$ | 19 | $0.86 \pm 2$ |
| 27 | Blue duiker | *Philantomba monticola* | 17 | $0.77 \pm 1.34$ | 16 | $0.72 \pm 1.24$ |
| 28 | Maxwell's duiker | *Philantomba maxwellii* | 16 | $0.72 \pm 1.3$ | 16 | $0.72 \pm 1.3$ |
| 29 | Egyptian goose | *Alopochen aegyptiaca* | 15 | $0.68 \pm 1.32$ | 15 | $0.68 \pm 1.32$ |
| 30 | Scimitar-horned oryx | *Oryx dammah* | 13 | $0.59 \pm 1$ | 13 | $0.59 \pm 1$ |
| 31 | Polar bear | *Ursus maritimus* | 12 | $0.54 \pm 0.96$ | 11 | $0.5 \pm 0.85$ |
| 32 | Aoudad | *Ammotragus lervia* | 12 | $0.54 \pm 0.73$ | 12 | $0.54 \pm 0.73$ |
| 33 | Olive baboon | *Papio anubis* | 11 | $0.5 \pm 0.8$ | 11 | $0.5 \pm 0.8$ |
| 34 | Spur-winged goose | *Plectropterus gambensis* | 10 | $0.45 \pm 0.85$ | 10 | $0.45 \pm 0.85$ |
| 35 | Cheetah | *Acinonyx jubatus* | 8 | $0.36 \pm 0.72$ | 7 | $0.36 \pm 0.56$ |
| 36 | Honey badger | *Mellivora capensis* | 8 | $0.36 \pm 0.65$ | 8 | $0.36 \pm 0.65$ |
| 37 | Markhor | *Capra falconeri* | 8 | $0.36 \pm 0.58$ | 8 | $0.36 \pm 0.58$ |
| 38 | Wild goat | *Capra hircus aegagrus* | 7 | $0.31 \pm 0.71$ | 7 | $0.31 \pm 0.71$ |
| 39 | Bighorn sheep | *Ovis canadensis* | 5 | $0.22 \pm 0.52$ | 5 | $0.22 \pm 0.52$ |
| 40 | American alligator | *Alligator mississippiensis* | 4 | $0.18 \pm 0.5$ | 4 | $0.18 \pm 0.5$ |
| 41 | Argali | *Ovis ammon* | 4 | $0.18 \pm 0.39$ | 4 | $0.18 \pm 0.39$ |
| 42 | Canada lynx | *Lynx canadensis* | 3 | $0.13 \pm 0.46$ | 3 | $0.13 \pm 0.46$ |
| 43 | Blesbok | *Damaliscus pygargus* | 3 | $0.13 \pm 0.46$ | 3 | $0.13 \pm 0.46$ |
| 44 | Gobi argali | *Ovis darwini* | 3 | $0.13 \pm 0.46$ | 3 | $0.13 \pm 0.46$ |
| 45 | Urial† | *Ovis aries* | 3 | $0.13 \pm 0.35$ | 3 | $0.13 \pm 0.35$ |
| 46 | Sitatunga | *Tragelaphus spekii* | 2 | $0.09 \pm 0.42$ | 2 | $0.09 \pm 0.42$ |
| 47 | Bay duiker | *Cephalophus dorsalis* | 2 | $0.09 \pm 0.42$ | 2 | $0.09 \pm 0.42$ |
| 48 | Blue sheep | *Pseudois nayaur* | 2 | $0.09 \pm 0.42$ | 2 | $0.09 \pm 0.42$ |
| 49 | Eurasian lynx | *Lynx lynx* | 2 | $0.09 \pm 0.29$ | 2 | $0.09 \pm 0.29$ |
| 50 | Bobcat | *Lynx rufus* | 2 | $0.09 \pm 0.29$ | 2 | $0.09 \pm 0.29$ |
| 51 | Walrus | *Odobenus rosmarus* | 2 | $0.09 \pm 0.29$ | 1 | $0.04 \pm 0.21$ |
| 52 | Western tur | *Capra caucasica* | 2 | $0.09 \pm 0.29$ | 2 | $0.09 \pm 0.29$ |
| 53 | Bongo | *Tragelaphus eurycerus* | 2 | $0.09 \pm 0.29$ | 2 | $0.09 \pm 0.29$ |
| 54 | Cape mountain zebra | *Equus zebra zebra* | 2 | $0.09 \pm 0.29$ | 2 | $0.09 \pm 0.29$ |
| 55 | Central African rock python | *Python sebae* | 2 | $0.09 \pm 0.29$ | 2 | $0.09 \pm 0.29$ |
| 56 | White-faced whistling duck | *Dendrocygna viduata* | 2 | $0.09 \pm 0.29$ | 2 | $0.09 \pm 0.29$ |
| 57 | Common jackal | *Canis aureus* | 1 | 0.04 ± 0.21 | 1 | 0.04 ± 0.21 |
| 58 | Striped hyena | *Hyaena hyaena* | 1 | 0.04 ± 0.21 | 1 | 0.04 ± 0.21 |
| 59 | North American river otter | *Lontra canadensis* | 1 | 0.04 ± 0.21 | 1 | 0.04 ± 0.21 |
| 60 | Aardwolf | *Proteles cristata* | 1 | 0.04 ± 0.21 | 1 | 0.04 ± 0.21 |
| 61 | Addax | *Addax nasomaculatus* | 1 | 0.04 ± 0.21 | 1 | 0.04 ± 0.21 |
| 62 | Hog deer | *Axis porcinus* | 1 | 0.04 ± 0.21 | 1 | 0.04 ± 0.21 |
| 63 | Wood bison | *Bison bison athabasca* | 1 | 0.04 ± 0.21 | 1 | 0.04 ± 0.21 |
| 64 | Nilgai | *Boselaphus tragocamelus* | 1 | 0.04 ± 0.21 | 1 | 0.04 ± 0.21 |
| 65 | Dama gazelle | *Nanger dama* | 1 | 0.04 ± 0.21 | 1 | 0.04 ± 0.21 |
| 66 | Arabian oryx | *Oryx leucoryx* | 1 | 0.04 ± 0.21 | 1 | 0.04 ± 0.21 |
| 67 | Collared peccary | *Pecari tajacu* | 1 | 0.04 ± 0.21 | 1 | 0.04 ± 0.21 |
| 68 | Blue monkey | *Cercopithecus mitis* | 1 | 0.04 ± 0.21 | 1 | 0.04 ± 0.21 |
| 69 | Guereza | *Colobus guereza* | 1 | 0.04 ± 0.21 | 1 | 0.04 ± 0.21 |
| 70 | Gelada baboon | *Theropithecus gelada* | 1 | 0.04 ± 0.21 | 1 | 0.04 ± 0.21 |
| 71 | Black rhino | *Diceros bicornis* | 1 | 0.04 ± 0.21 | 1 | 0.04 ± 0.21 |
| 72 | Bald eagle | *Haliaeetus leucocephalus* | 1 | 0.04 ± 0.21 | 1 | 0.04 ± 0.21 |
| 73 | Maro Polo argali | *Ovis polii* | 1 | 0.04 ± 0.21 | 1 | 0.04 ± 0.21 |

\*For lion, trophies sourced from the wild totaled 89 (4.04 ± 6.03 trophies/year; 3.54 ± 5.5 WOE/year), trophies sourced from animals bred in captivity totaled 59 (8.42 ± 2.43 trophies/year; 8.42 ± 2.43 WOE/year) and trophies sourced from animals born in captivity totaled 1 (0.04 ± .021 trophies/year; 0.04 ± .021 WOE/year).
†We include *Ovis aries* in this table but in table S12 refer to two species: mouflon (*O. gmelini*) and urial (*O. vignei*) following taxonomic updates on the IUCN Red List.

**Table S9.**
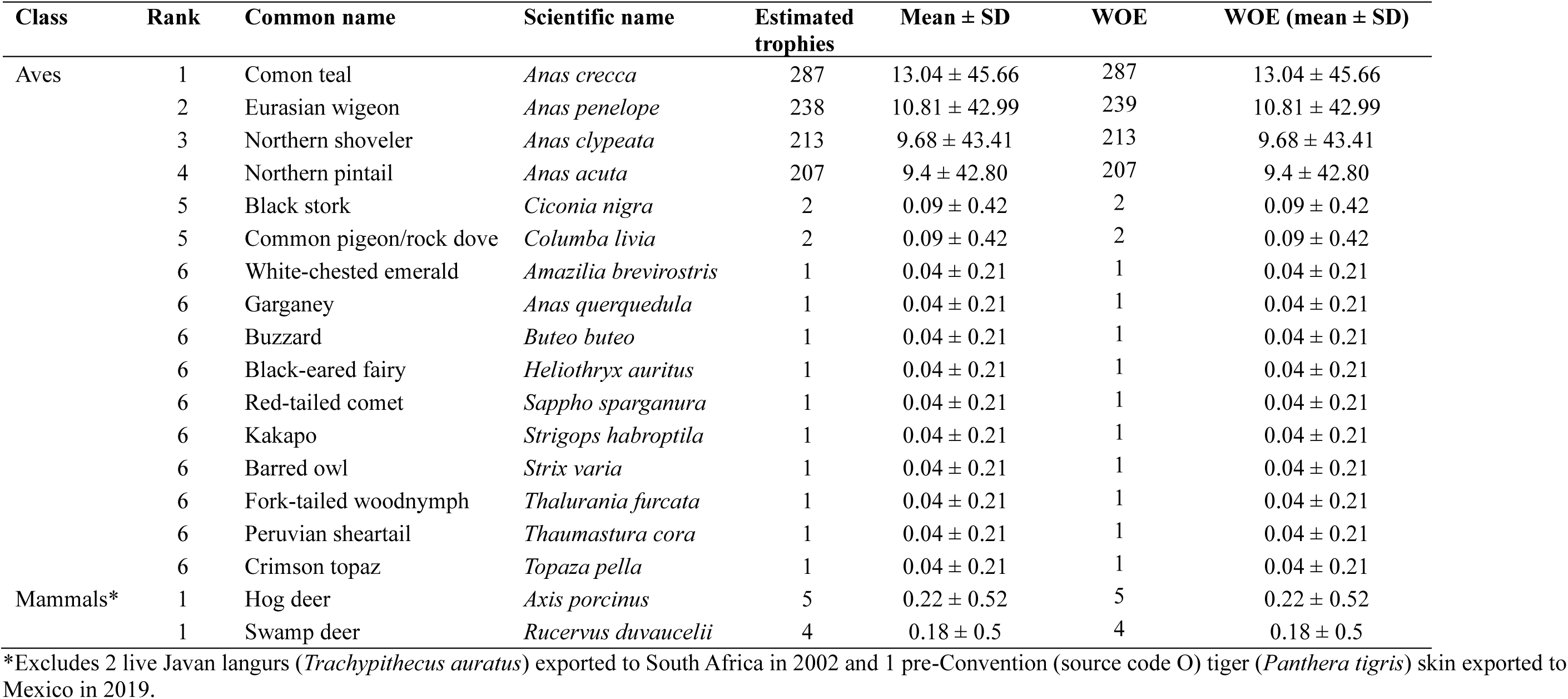
Species exported (n=18) from the UK as hunting trophies (2000-2021), estimated number of trophies, mean (± SD) number of trophies traded/year, WOEs traded in this period, and mean (± SD) WOEs traded in this period/year, ranked by number of trophies.

| Class | Rank | Common name | Scientific name | Estimated trophies | Mean ± SD | WOE | WOE (mean ± SD) |
| --- | --- | --- | --- | --- | --- | --- | --- |
| Aves | 1 | Comon teal | *Anas crecca* | 287 | 13.04 ± 45.66 | 287 | 13.04 ± 45.66 |
| | 2 | Eurasian wigeon | *Anas penelope* | 238 | 10.81 ± 42.99 | 239 | 10.81 ± 42.99 |
| | 3 | Northern shoveler | *Anas clypeata* | 213 | 9.68 ± 43.41 | 213 | 9.68 ± 43.41 |
| | 4 | Northern pintail | *Anas acuta* | 207 | 9.4 ± 42.80 | 207 | 9.4 ± 42.80 |
| | 5 | Black stork | *Ciconia nigra* | 2 | 0.09 ± 0.42 | 2 | 0.09 ± 0.42 |
| | 5 | Common pigeon/rock dove | *Columba livia* | 2 | 0.09 ± 0.42 | 2 | 0.09 ± 0.42 |
| | 6 | White-chested emerald | *Amazilia brevirostris* | 1 | 0.04 ± 0.21 | 1 | 0.04 ± 0.21 |
| | 6 | Garganey | *Anas querquedula* | 1 | 0.04 ± 0.21 | 1 | 0.04 ± 0.21 |
| | 6 | Buzzard | *Buteo buteo* | 1 | 0.04 ± 0.21 | 1 | 0.04 ± 0.21 |
| | 6 | Black-eared fairy | *Heliothryx auritus* | 1 | 0.04 ± 0.21 | 1 | 0.04 ± 0.21 |
| | 6 | Red-tailed comet | *Sappho sparganura* | 1 | 0.04 ± 0.21 | 1 | 0.04 ± 0.21 |
| | 6 | Kakapo | *Strigops habroptila* | 1 | 0.04 ± 0.21 | 1 | 0.04 ± 0.21 |
| | 6 | Barred owl | *Strix varia* | 1 | 0.04 ± 0.21 | 1 | 0.04 ± 0.21 |
| | 6 | Fork-tailed woodnymph | *Thalurania furcata* | 1 | 0.04 ± 0.21 | 1 | 0.04 ± 0.21 |
| | 6 | Peruvian sheartail | *Thaumastura cora* | 1 | 0.04 ± 0.21 | 1 | 0.04 ± 0.21 |
| | 6 | Crimson topaz | *Topaza pella* | 1 | 0.04 ± 0.21 | 1 | 0.04 ± 0.21 |
| Mammals* | 1 | Hog deer | *Axis porcinus* | 5 | 0.22 ± 0.52 | 5 | 0.22 ± 0.52 |
| | 1 | Swamp deer | *Rucervus duvaucelii* | 4 | 0.18 ± 0.5 | 4 | 0.18 ± 0.5 |

\*Excludes 2 live Javan langurs (*Trachypithecus auratus*) exported to South Africa in 2002 and 1 pre-Convention (source code O) tiger (*Panthera tigris*) skin exported to Mexico in 2019.

N.B. Exports of hunting trophies from the UK involved only 17 trophies in 2015-2021. They have not been tabulated but included the following: 5 *A. porcinus* trophies (as per Table S9), 4 of which were from captive sources (source code C) and 1 was a previously seized trophy (source code I); 3 *R. duvaucelli* trophies all from captive sources (source code C), 2 trophies from *C. nigra* (source code W) and single trophies from *A. brevirostris*, *H. auratus*, *S. sparganura*, *S. habroptila*, *T. cora, T. furcate*, and *T. pella* (source code O, pre-Convention) as per Table S9.

**Table S10.**
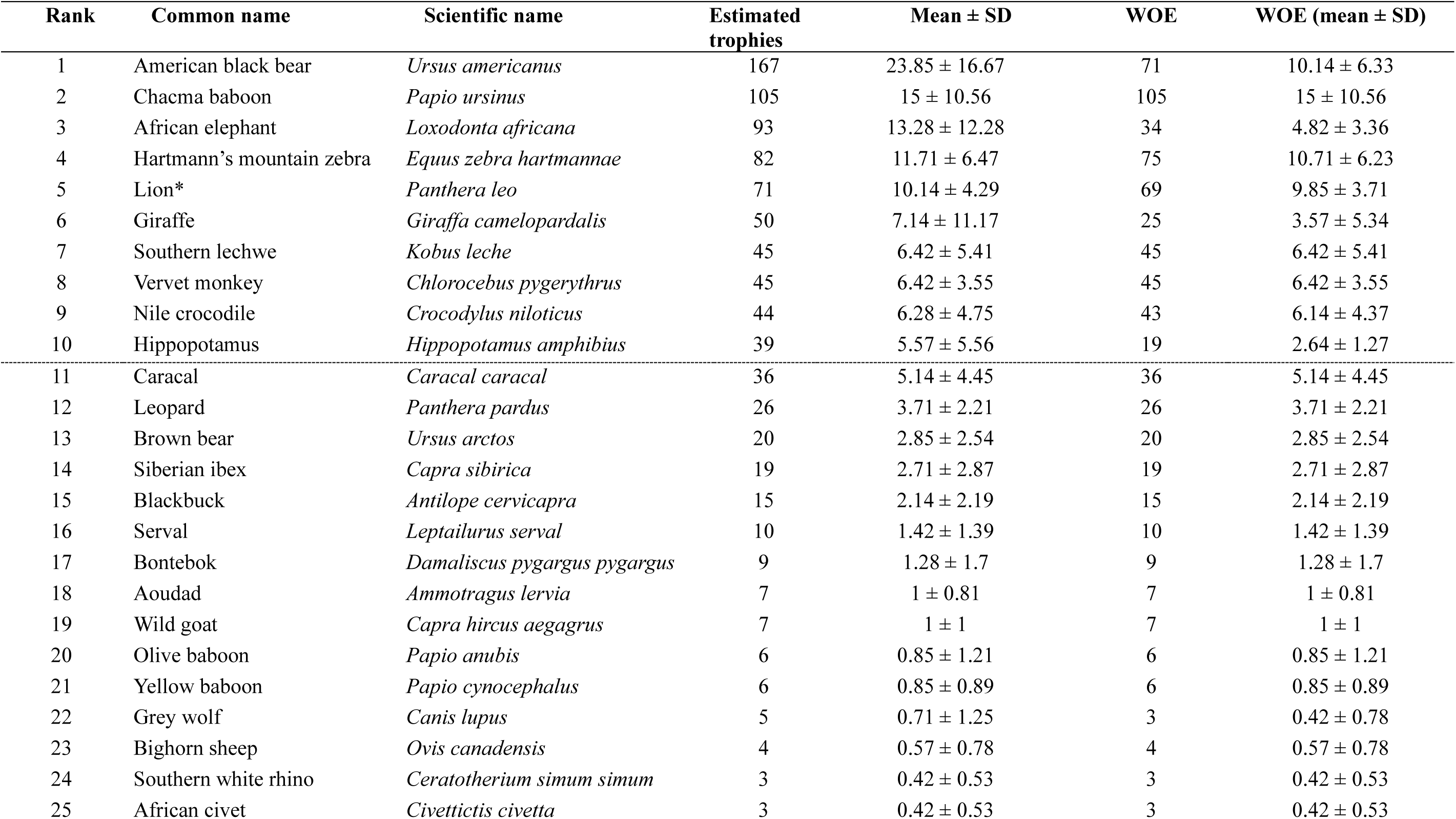
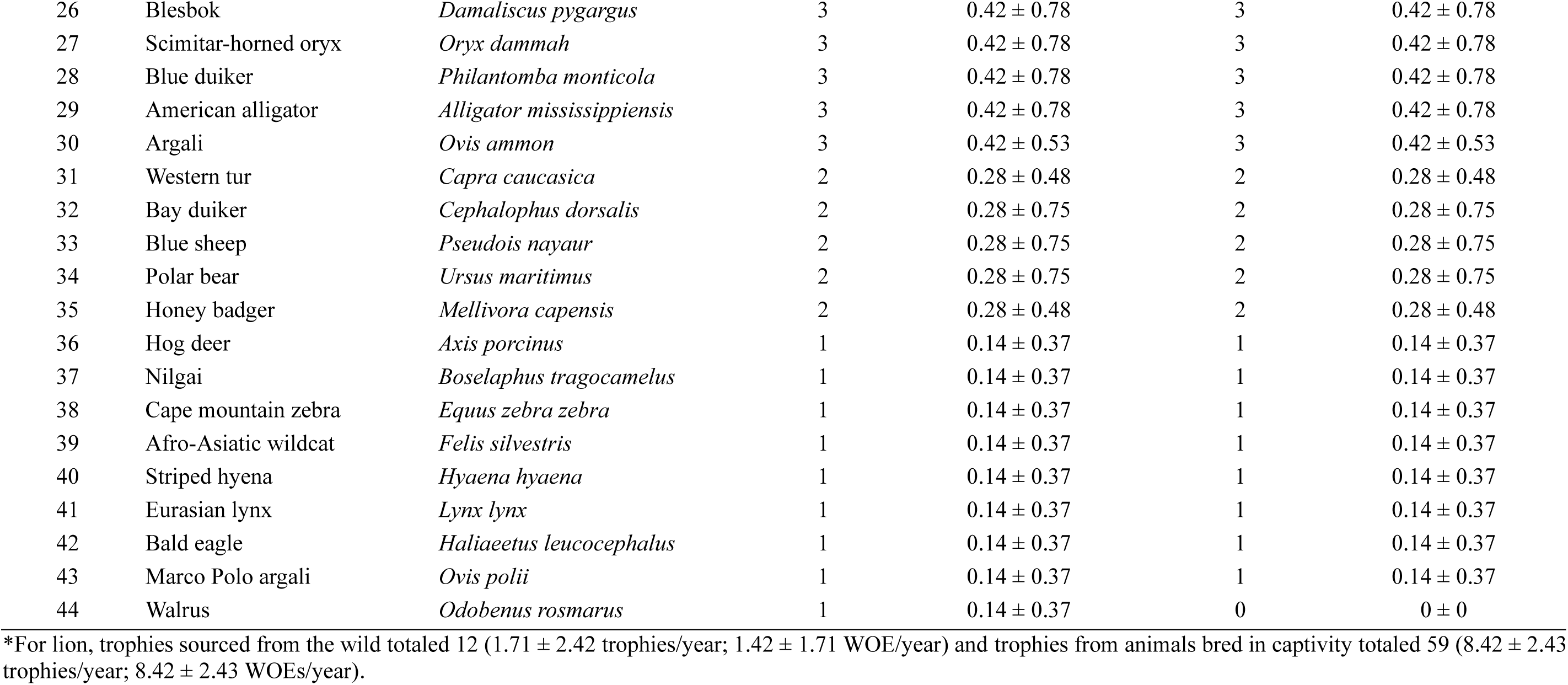
Species and subspecies imported (n=44) to the UK as hunting trophies (2015-2021), estimated number of trophies, mean (± SD) number of trophies traded/year, WOEs traded in this period, and mean (± SD) WOEs traded in this period/year, ranked by number of trophies.

### Supplementary Material 3 – Results – Population status

**Table S11.** Population status of species and subspecies exported to the UK as hunting trophies (2015-2021) in countries granting the export(s). Species ranked by number of trophies. AR=Argentina, AU=Australia, AZ=Azerbaijan, BJ=Benin, BW=Botswana, CM=Cameroon, CA=Canada, CG=Congo, KG=Kyrgyzstan, MX=Mexico, MZ=Mozambique, NA=Namibia, NP=Nepal, PK=Pakistan, RU=Russian Federation, TR=Türkiye, TZ= Tanzania, UG=Uganda, US=United States of America, ZA=South Africa, ZM=Zambia, ZW=Zimbabwe. Except for CITES (2019, 2022), Giraffe Conservation Foundation (2019) and Mech & Boitani (2004) sources are global or regional Red List assessments.

| Common name | Scientific name | Exporter | No. of trophies | Population status (e.g., increasing, decreasing, stable or unknown) | Is population stable, increasing or similar? | Source |
| --- | --- | --- | --- | --- | --- | --- |
| American black bear | *Ursus americanus* | CA | 167 | Increasing (estimated at 450,000 individuals) | Yes | Garshelis *et al.* 2016 |
| Chacma baboon | *Papio ursinus* | ZA | 70 | Population declining globally but common and widespread | Yes | Sithaldeen 2019 |
|  |  | NA | 20 | Population declining globally but common and widespread | Yes | Sithaldeen 2019 |
|  |  | ZW | 14 | Population declining globally but common and widespread | Yes | Sithaldeen 2019 |
|  |  | ZM | 1 | Population declining globally but common and widespread | Yes | Sithaldeen 2019 |
| African elephant | *Loxodonta africana* | ZW | 60 | Abundant, population estimated at 82,630 individuals | Yes | CITES 2022 |
|  |  | ZA | 15 | Abundant, population estimated at 26,864 individuals and increasing | Yes | Selier *et al.* 2016 |
|  |  | BW | 5 | Abundant, population estimated at 131,626 individuals | Yes | CITES 2022 |
|  |  | MZ | 5 | Unknown | Unknown |  |
|  |  | NA | 4 | Abundant, population estimated at 22,754 individuals | Yes | CITES 2022 |
|  |  | TZ | 2 | Unknown | Unknown |  |
|  |  | ZM | 2 | Stable, population estimated at 7,000 individuals | Yes | CITES 2019 |
| Hartmann's Mountain zebra | *Equus zebra hartmannae* | NA | 71 | Increasing | Yes | Gosling *et al.* 2019 |
|  |  | ZA | 11 | Increasing | Yes | Gosling *et al.* 2019 |
| Lion | *Panthera leo* | ZA | 61 | Increasing | Yes | Bauer *et al.* 2016 |
|  |  | NA | 6 | Increasing† | Yes | Bauer *et al.* 2016 |
|  |  | MZ | 2 | Unknown | Unknown | Bauer *et al.* 2016 |
|  |  | TZ | 2 | Declining | No | Bauer *et al.* 2016 |
| Giraffe | *Giraffa giraffa giraffa* | ZA | 43 | Population increased in recent decades, now numbers 37,000 individuals | Yes | Giraffe Conservation Foundation (2019) |
|  |  | ZW | 5 | Population increased in recent decades, now numbers 37,000 individuals | Yes | Giraffe Conservation Foundation (2019) |
|  | *Giraffa giraffa angolensis* | NA | 2 | Population increased in recent decades, now numbers 17,750 individuals | Yes | Giraffe Conservation Foundation (2019) |
| Southern lechwe | *Kobus leche* | ZA | 34 | Extra-limital population | Extra-limital | IUCN SSC Antelope Specialist Group 2017a |
|  |  | NA | 6 | Some subspecies populations increasing, other declining | Unknown | IUCN SSC Antelope Specialist Group 2017a |
|  |  | ZM | 5 | Some subspecies populations increasing, other declining | Unknown | IUCN SSC Antelope Specialist Group 2017a |
| Vervet monkey | *Chlorocebus pygerythrus* | ZA | 41 | Global population declining but locally abundant | Unknown | Butynski *et al.* 2022 |
|  |  | ZM | 2 | Global population declining but locally abundant | Unknown | Butynski *et al.* 2022 |
|  |  | ZW | 2 | Global population declining but locally abundant | Unknown | Butynski *et al.* 2022 |
| Nile crocodile | *Crocodylus niloticus* | ZA | 33 | Stable | Yes | Turner *et al.* 2017 |
|  |  | ZW | 5 | Increasing in places, declining in others | No | Isberg *et al.* 2019 |
|  |  | MZ | 2 | Unknown | Unknown | Isberg *et al.* 2019 |
|  |  | NA | 2 | Increasing in places, unknown in others | Unknown | Isberg *et al.* 2019 |
|  |  | TZ | 2 | Unknown | Unknown | Isberg *et al.* 2019 |
| Hippopotamus | *Hippopotamus amphibius* | ZW | 15 | Stable | Yes | Lewison & Pluháček 2017 |
|  |  | ZM | 12 | Stable | Yes | Lewison & Pluháček 2017 |
|  |  | ZA | 8 | Stable in southern Africa and most subpopulations in South Africa have increased in the last decade | Yes | Eksteen *et al.* 2016 |
|  |  | TZ | 4 | Unknown, but (dated) population estimate is 20,000 individuals and considered and underestimate | Unknown | Lewison & Pluháček 2017 |
| Caracal | *Caracal caracal* | ZA | 34 | Common and expanding into new areas | Yes | Avgan *et al.* 2016 |
|  |  | NA | 2 | Common and expanding into new areas | Yes | Avgan *et al.* 2016 |
| Leopard | *Panthera pardus* | MZ | 4 | Declining | No | Stein *et al.* 2020 |
|  |  | NA | 8 | Unknown | No | Stein *et al.* 2020 |
|  |  | TZ | 7 | Unknown | No | Stein *et al.* 2020 |
|  |  | ZA | 1 | Declining | No | Stein *et al.* 2020, Swanepoel *et al.* 2016a |
|  |  | ZM | 2 | Declining | No | Stein *et al.* 2020 |
|  |  | ZW | 4 | Declining | No | Stein *et al.* 2020 |
| Brown bear | *Ursus arctos* | RU | 16 | Abundant, population estimated at 100,000 individuals | Yes | McLellan *et al.* 2017 |
|  |  | US | 4 | Abundant, population estimated at 33,000 individuals | Yes | McLellan *et al.* 2017 |
| Siberian ibex | *Capra sibirica* | PK | 1 | Locally abundant in the north of the country | Unknown | Reading *et al.* 2020a |
|  |  | KG | 18 | Abundant, population estimated at 70,000 individuals (includes Tajikistan) | Yes | Reading *et al.* 2020a |
| Blackbuck | *Antelope cervicapra* | AR | 14 | Extra-limital population | Extra-limital | IUCN SSC Antelope Specialist Group 2017b |
|  |  | US | 1 | Extra-limital population | Extra-limital | IUCN SSC Antelope Specialist Group 2017b |
| Serval | *Leptailurus serval* | TZ | 2 | Unknown | Unknown | Thiel 2019 |
|  |  | ZA | 8 | Stable | Yes | Ramesh *et al.* 2016 |
| Bontebok | *Damaliscus pygargus pygargus* | ZA | 9 | Stable (some subpopulations increasing) | Yes | Radloff *et al.* 2016, 2017 |
| Aoudad | *Ammotragus lervia* | US | 2 | Extra-limital population | Extra-limital | Cassinello *et al.* 2022 |
|  |  | ZA | 5 | Extra-limital population | Extra-limital | Cassinello *et al.* 2022 |
| Wild goat | *Capra hircus aegagrus* | TR | 7 | Declining (though might have stabilised) | No | Weinberg & Ambarli 2020 |
| Olive baboon | *Papio anubis* | BJ | 1 | Global population stable, widespread and locally common | Yes | Wallis 2020a |
|  |  | CM | 2 | Global population stable, widespread and locally common | Yes | Wallis 2020a |
|  |  | TZ | 2 | Global population stable, widespread and locally common | Yes | Wallis 2020a |
|  |  | UG | 1 | Global population stable, widespread and locally common | Yes | Wallis 2020a |
| Yellow baboon | *Papio cynocephalus* | TZ | 1 | Global population stable, widespread and locally common | Yes | Wallis 2020b |
|  |  | ZM | 5 | Global population stable, widespread and locally common | Yes | Wallis 2020b |
| Grey wolf | *Canis lupus* | CA | 5 | Stable (population estimated at c.40,000 individuals) | Yes | Boitani *et al.* 2018, Mech & Boitani 2004 |
| Bighorn sheep | *Ovis canadensis* | MX | 4 | Population estimated at 12,655-14,000 individuals and is increasing in many areas | Yes | Festa-Bianchet 2020 |
| Southern white rhino | *Ceratotherium simum simum* | ZA | 3 | Population declining globally but an estimated 18,064 individuals in South Africa | Yes | Emslie 2020a |
| African civet | *Civettictis civetta* | ZA | 2 | Population estimates suggest healthy populations | Yes | Swanepoel *et al.* 2016b |
|  |  | ZW | 1 | Unknown | Unknown | - |
| Blesbok | *Damaliscus pygargus* | ZA | 3 | Increasing | Yes | Dalton *et al.* 2019 |
| Scimitar-horned oryx | *Oryx dammah* | ZA | 3 | Extra-limital population | Extra-limital | IUCN SSC Antelope Specialist Group 2016b |
| Blue duiker | *Philantomba monticola* | ZA | 3 | Declining | No | Venter *et al.* 2016 |
| American alligator | *Alligator mississippiensis* | US | 3 | Increasing | Yes | Elsey *et al.* 2019 |
| Argali | *Ovis ammon* | KG | 3 | Population estimated at 16,669 individuals but abundance varies greatly in country | Unknown | Reading *et al.* 2020b |
| Western tur | *Capra caucasica* | AZ | 1 | Unknown | Unknown | - |
|  |  | RU | 1 | Unknown | Unknown | - |
| Bay duiker | *Cephalophus dorsalis* | CG | 2 | Declining (though stable in places) | No | IUCN SSC Antelope Specialist Group 2020 |
| Blue sheep | *Pseudois nayaur* | NP | 2 | Population in Nepal possibly 1947-10,000 individuals but this is uncertain | No | Harris 2014 |
| Polar bear | *Ursus maritimus* | CA | 2 | Unknown | Unknown | Wiig *et al.* 2015 |
| Honey badger | *Mellivora capensis* | ZA | 2 | Stable or increasing | Yes | Begg *et al.* 2016 |
| Hog deer | *Axis porcinus* | AU | 1 | Trophies were from captive animals | Extra-limital | - |
| Nilgai | *Boselaphus tragocamelus* | US | 1 | Trophies were from captive animals | Extra-limital | - |
| Cape Mountain zebra | *Equus zebra zebra* | ZA | 1 | Increasing | Yes | Hrabar *et al.* 2019 |
| Afro-Asiatic wildcat | *Felis silvestris* | ZA | 1 | Stable (but possibly declining) | Yes | Herbst *et al.* 2016 |
| Striped hyena | *Hyaena hyaena* | TZ | 1 | Unknown | No | AbiSaid and Dloniak 2015 |
| Eurasian lynx | *Lynx lynx* | RU | 1 | Stable in places, declining in others | No | Breitenmoser *et al.* 2015 |
| Walrus | *Odobenus rosmarus* | CA | 1 | Global population likely >225,000 but Canadian population unknown | Unknown | Lowry 2016 |
| Marco Polo argali | *Ovis polii* | KG | 1 | Population estimated at 4013 individuals but abundance varies greatly in country | Unknown | Reading *et al.* 2020b |
| Bald eagle | *Haliaeetus leucocephalus* | CA | 1 | Increasing | Yes | BirdLife International 2016 |

### Supplementary Material 4 – Results – Trophy hunting as a contributor to species being of elevated conservation concern

**Table S12.**
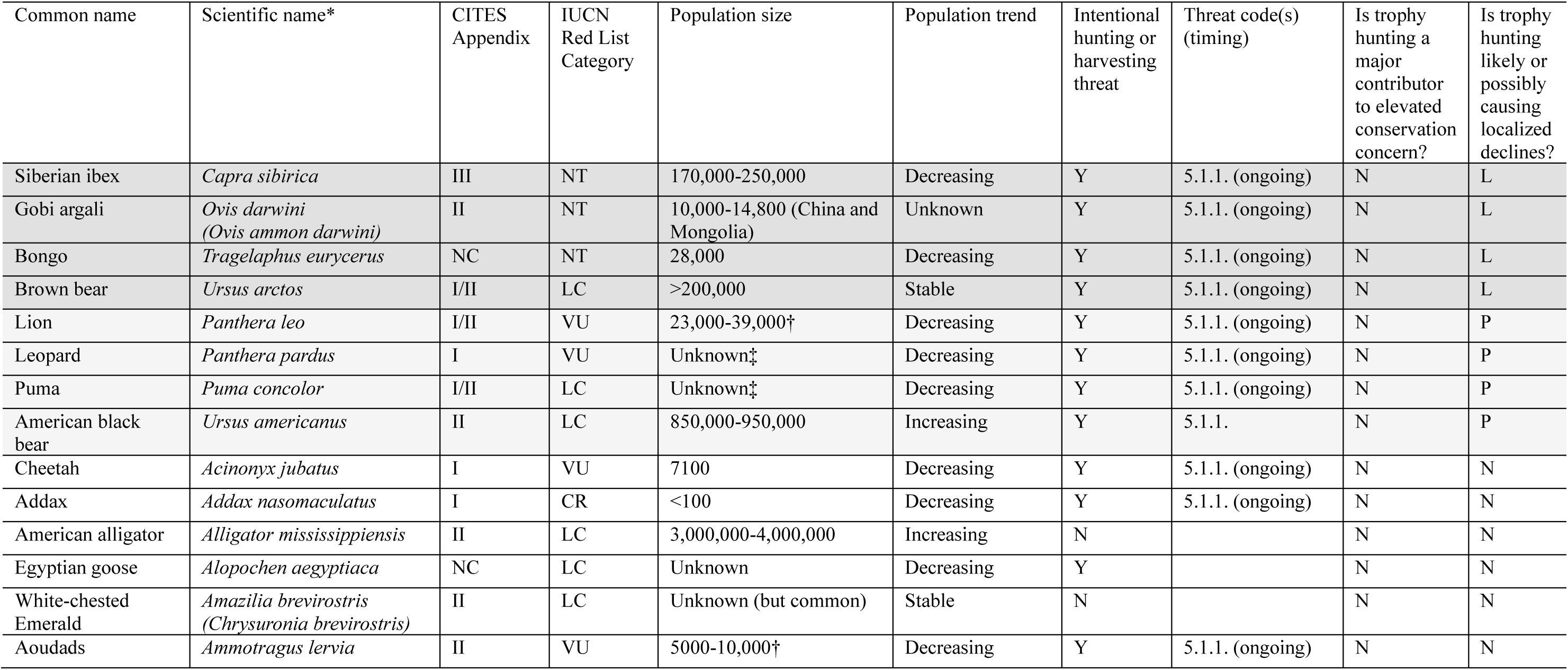
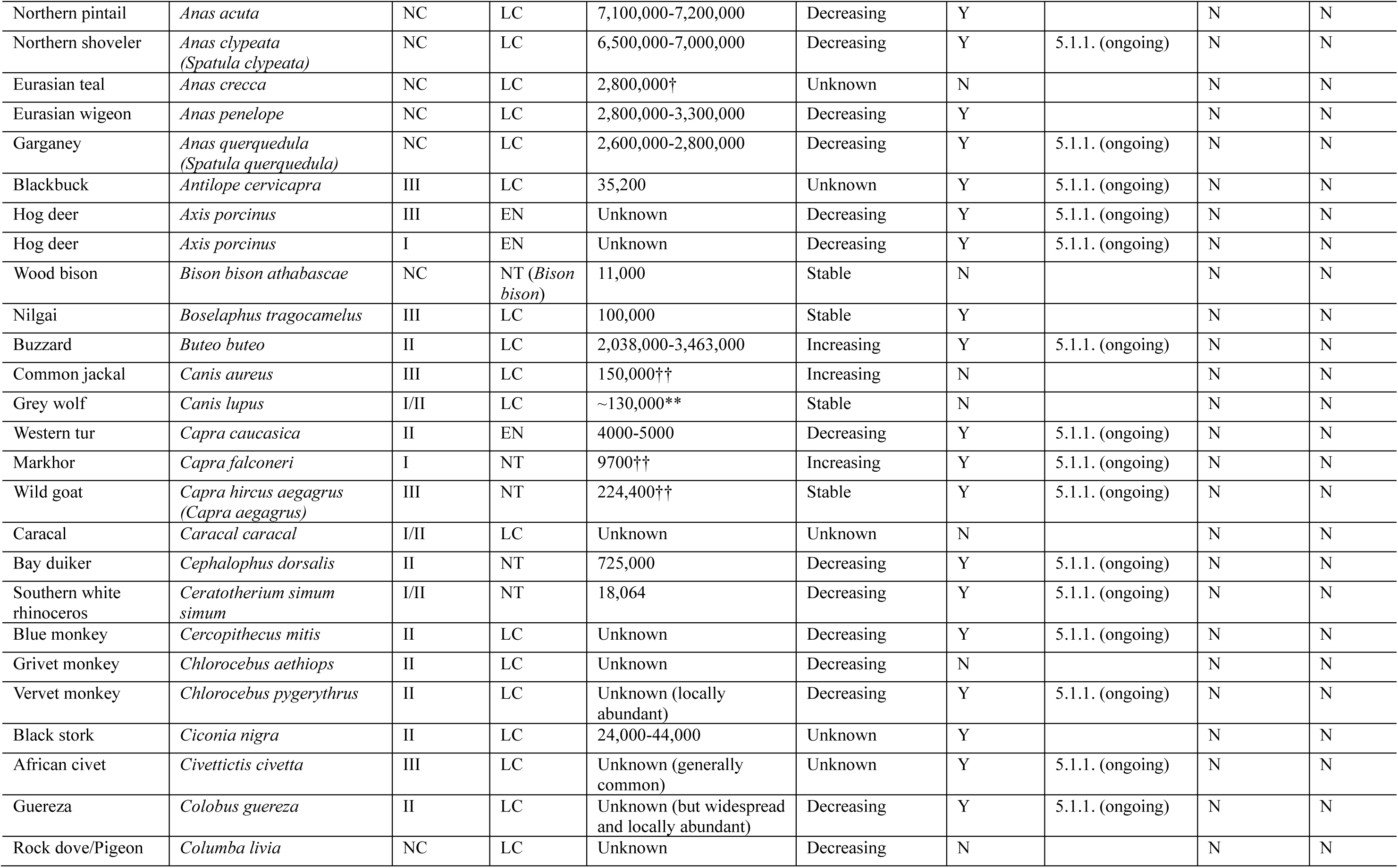
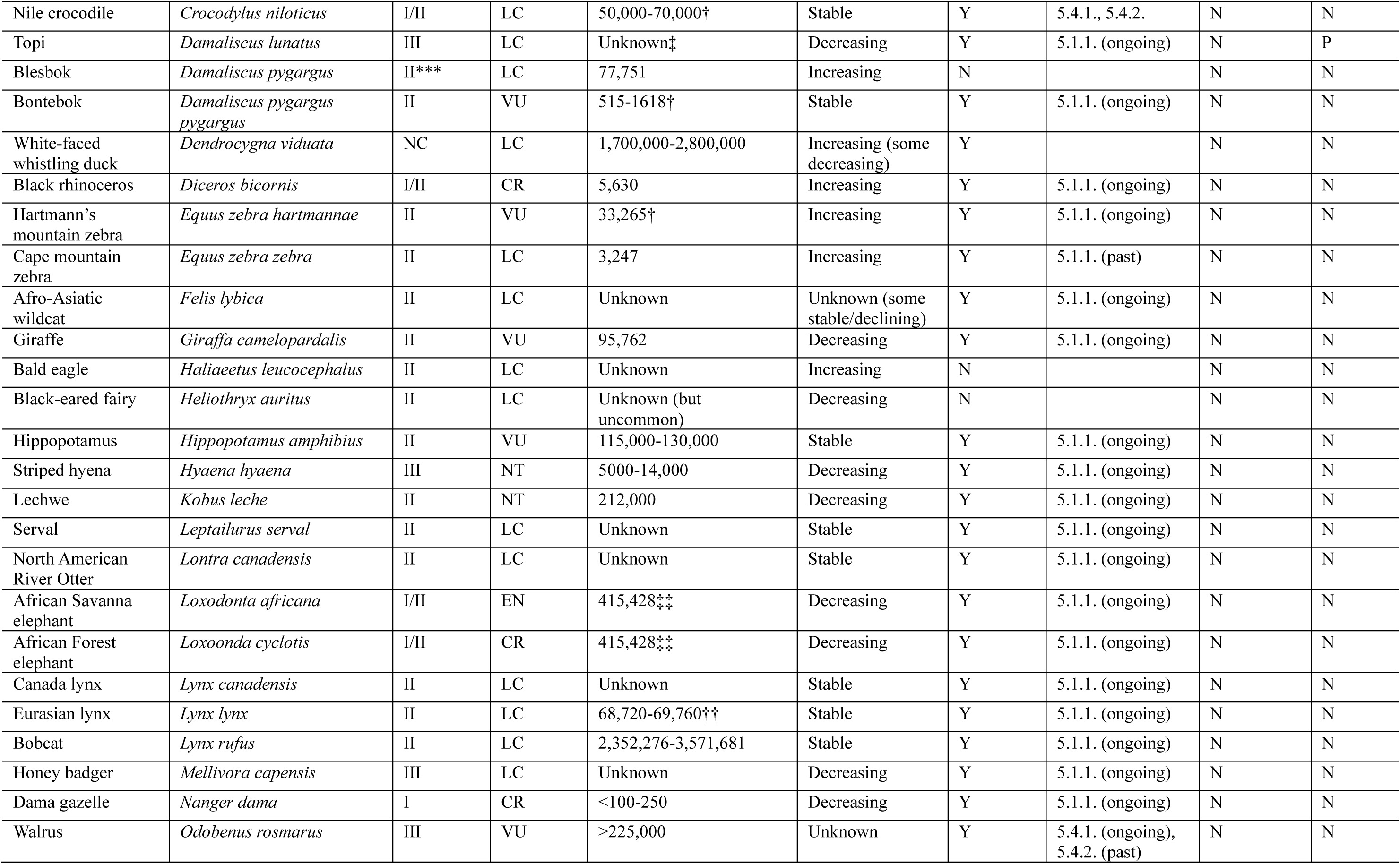
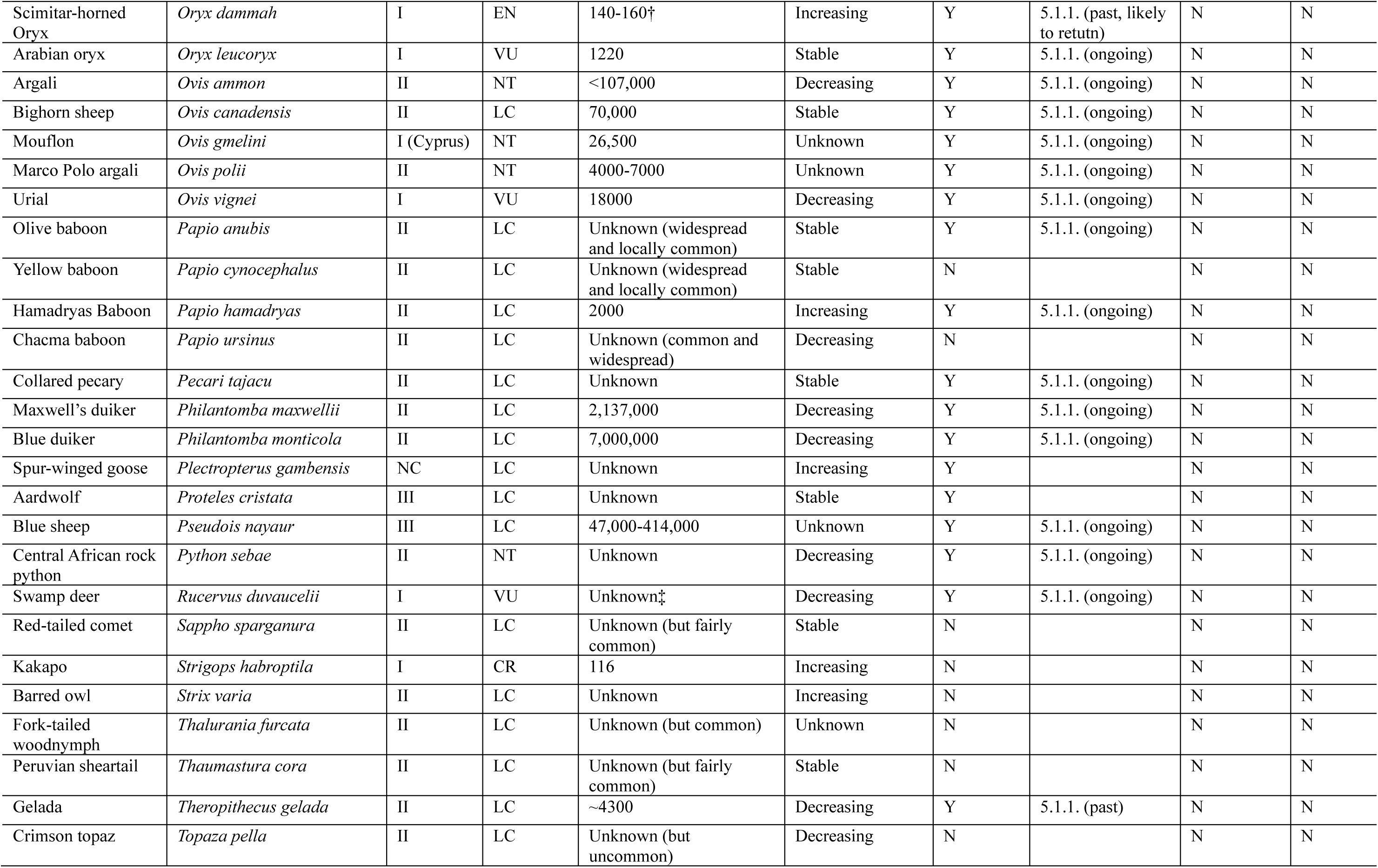
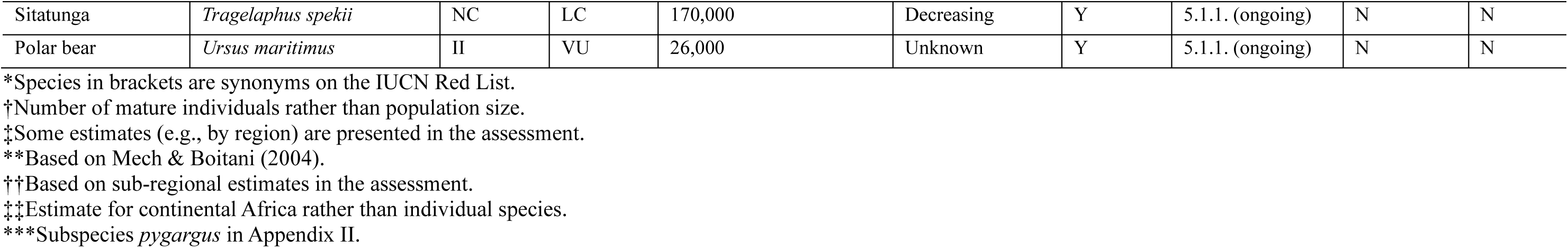
Species and subspecies imported to/exported from the UK as hunting trophies (2000-2021), CITES Appendix, IUCN Red List Threat Category, population size, population trend, and information on intentional hunting/harvesting threats, including whether trophy hunting poses a threat to species at different levels, or not. This table includes 93 species because the Red List treats the urialas mouflon (*O. gmelini*) and urial (*O. vignei*), and African elephant (*Loxodonta africana*) as the African Savanna Elephant (*L. africana*) and African Forest Elephant (*L. cyclotis*) following taxonomic updates.

**Table S13.** Species and subspecies for which trophy hunting provides, or has the potential to provide, benefits – to species and/or people – based on information in the threats, use and trade, and conservation actions section of Red List assessments.

| Common name | Scientific name |
| --- | --- |
| Aoudad | *Ammotragus lervia* |
| Markhor | *Capra falconeri* |
| Siberian ibex | *Capra sibirica* |
| White rhino | *Ceratotherium simum* |
| Blesbok | *Damaliscus pygargus* |
| Bontebok | *Damaliscus pygargus pygargus* |
| Black rhino | *Diceros bicornis* |
| Hartmann's Mountain zebra | *Equus zebra hartmannae* |
| Cape Mountain zebra | *Equus zebra zebra* |
| Southern lechwe | *Kobus leche* |
| Argali | *Ovis ammon* |
| Bighorn sheep` | *Ovis canadensis* |
| Gobi argali | *Ovis darwini* |
| Marco Polo argali | *Ovis polii* |
| Lion | *Panthera leo* |
| Leopard | *Panthera pardus* |
| Blue sheep | *Pseudois nayaur* |
| Bongo | *Tragelaphus eurycerus* |
| Sitatunga | *Tragelaphus spekii* |
| American black bear | *Ursus americanus* |

### Supp. Material 5 – Results – Evaluating the UK Government’s impact assessment of the bill

**Table S14.**
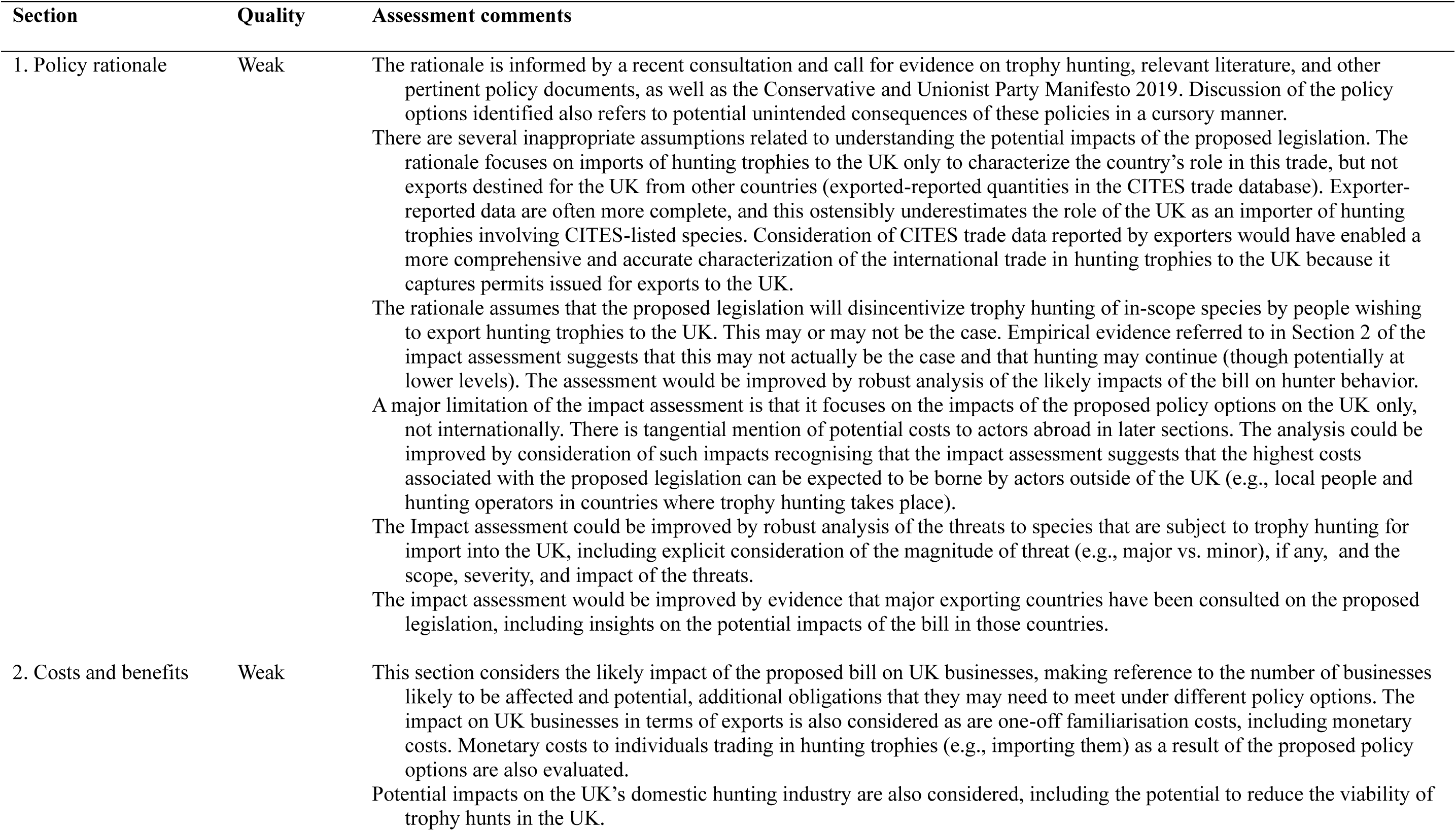
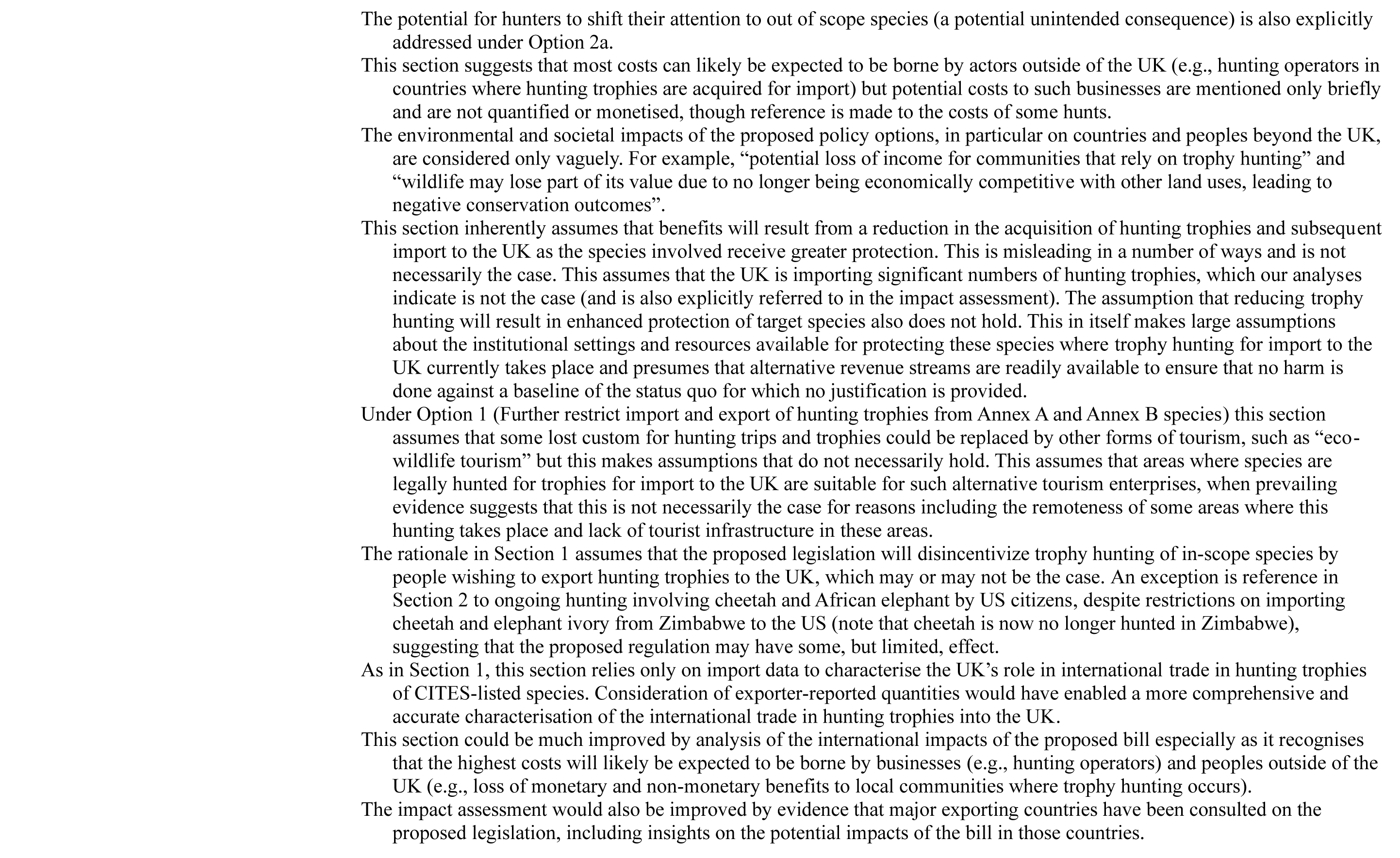
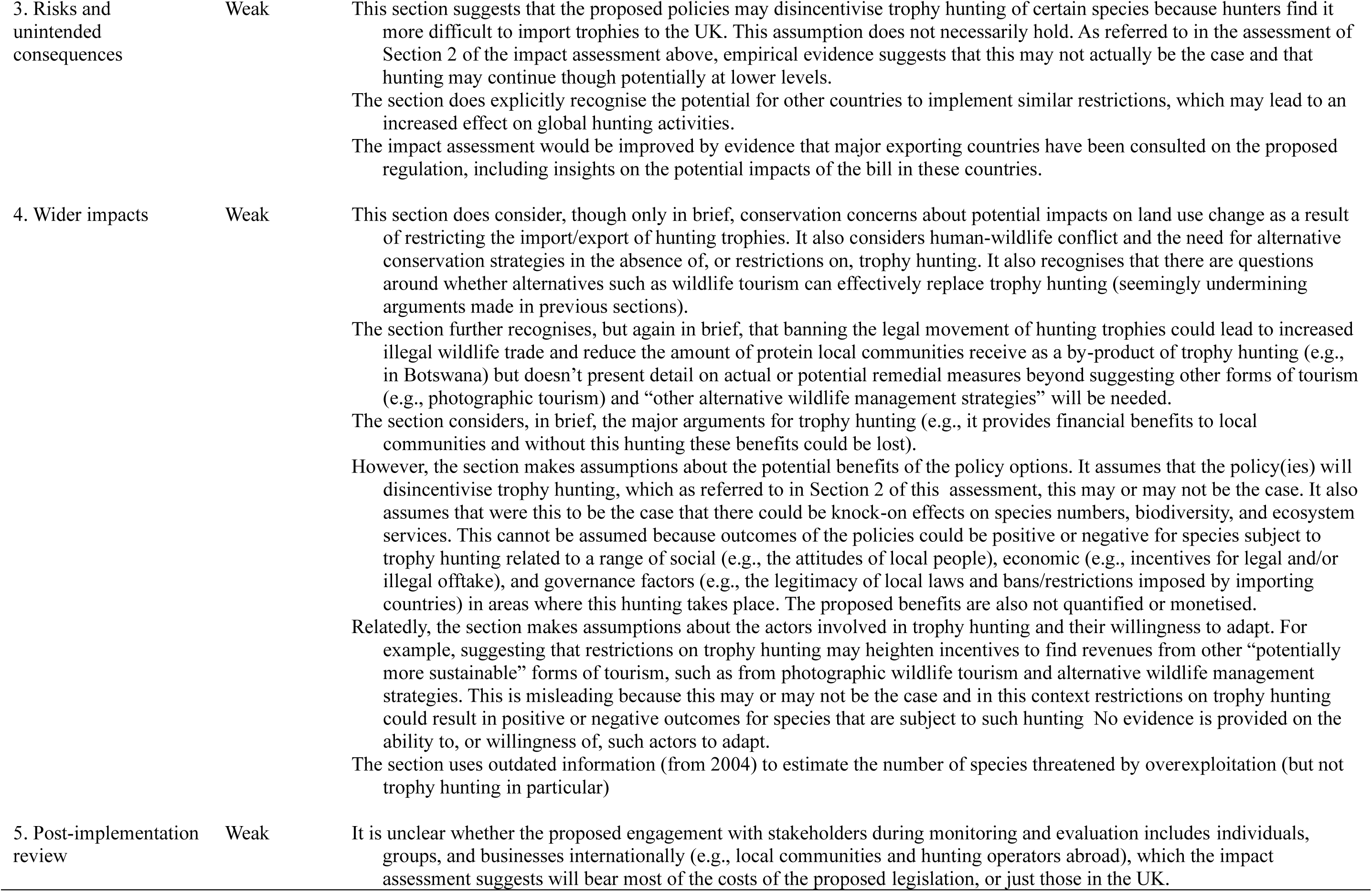
Assigned quality indicator and associated comments for the Hunting Trophies (Import Prohibition) Bill impact assessment.

### Supplementary Material 6 – Discussion

**Table S15.**
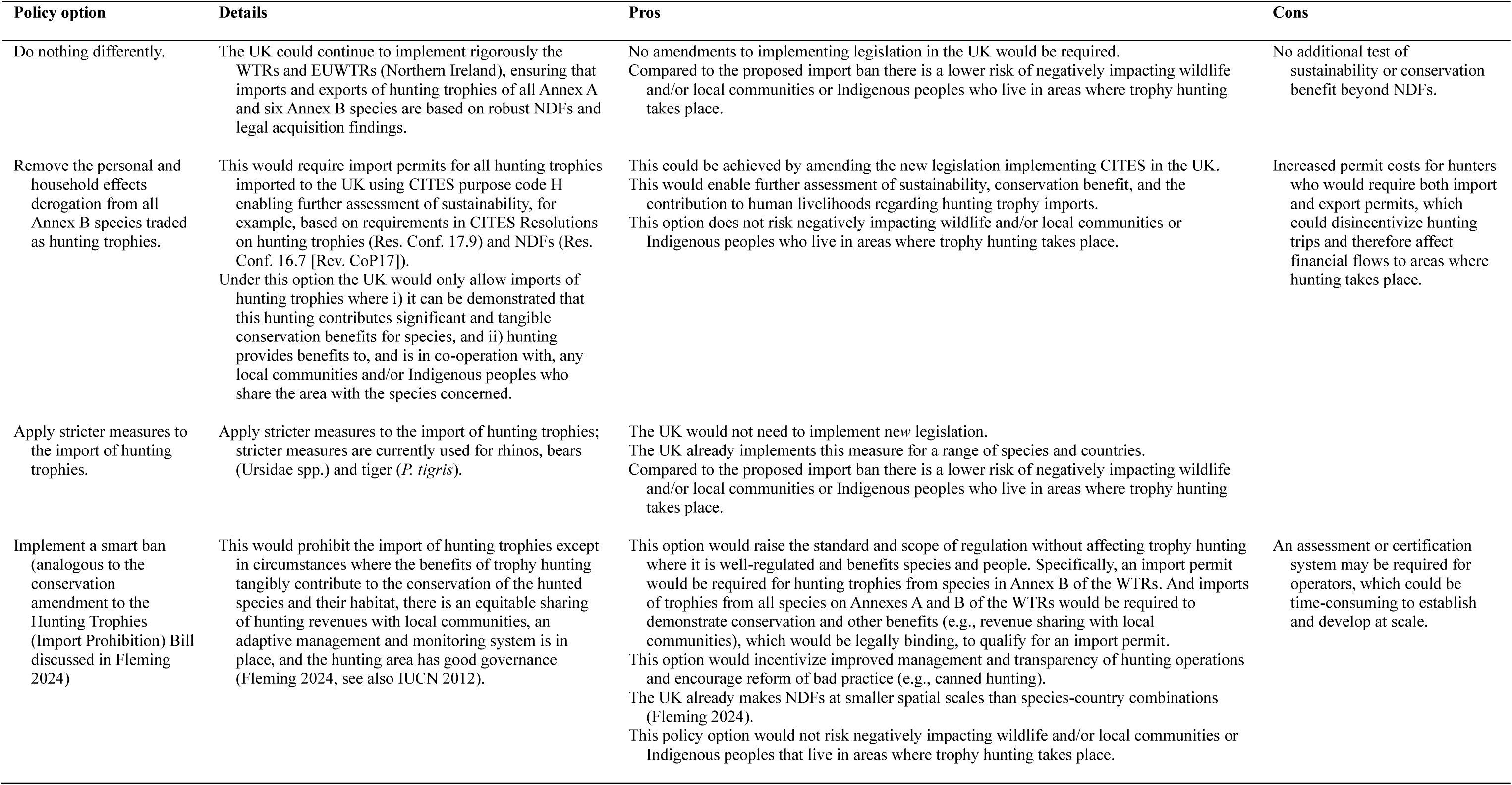
Non-exhaustive pros and cons of alternative regulatory options.

| Policy option | Details | Pros | Cons |
| --- | --- | --- | --- |
| Do nothing differently. | The UK could continue to implement rigorously the WTRs and EUWTRs (Northern Ireland), ensuring that imports and exports of hunting trophies of all Annex A and six Annex B species are based on robust NDFs and legal acquisition findings. | No amendments to implementing legislation in the UK would be required. Compared to the proposed import ban there is a lower risk of negatively impacting wildlife and/or local communities or Indigenous peoples who live in areas where trophy hunting takes place. | No additional test of sustainability or conservation benefit beyond NDFs. |
| Remove the personal and household effects derogation from all Annex B species traded as hunting trophies. | This would require import permits for all hunting trophies imported to the UK using CITES purpose code H enabling further assessment of sustainability, for example, based on requirements in CITES Resolutions on hunting trophies (Res. Conf. 17.9) and NDFs (Res. Conf. 16.7 [Rev. CoP17]).<br>Under this option the UK would only allow imports of hunting trophies where i) it can be demonstrated that this hunting contributes significant and tangible conservation benefits for species, and ii) hunting provides benefits to, and is in co-operation with, any local communities and/or Indigenous peoples who share the area with the species concerned. | This could be achieved by amending the new legislation implementing CITES in the UK. This would enable further assessment of sustainability, conservation benefit, and the contribution to human livelihoods regarding hunting trophy imports. This option does not risk negatively impacting wildlife and/or local communities or Indigenous peoples who live in areas where trophy hunting takes place. | Increased permit costs for hunters who would require both import and export permits, which could disincentivize hunting trips and therefore affect financial flows to areas where hunting takes place. |
| Apply stricter measures to the import of hunting trophies. | Apply stricter measures to the import of hunting trophies; stricter measures are currently used for rhinos, bears (*Ursidae* spp.) and tiger (*P. tigris*). | The UK would not need to implement new legislation. The UK already implements this measure for a range of species and countries. Compared to the proposed import ban there is a lower risk of negatively impacting wildlife and/or local communities or Indigenous peoples who live in areas where trophy hunting takes place. |  |
| Implement a smart ban (analogous to the conservation amendment to the Hunting Trophies (Import Prohibition) Bill discussed in Fleming 2024) | This would prohibit the import of hunting trophies except in circumstances where the benefits of trophy hunting tangibly contribute to the conservation of the hunted species and their habitat, there is an equitable sharing of hunting revenues with local communities, an adaptive management and monitoring system is in place, and the hunting area has good governance (Fleming 2024, see also IUCN 2012). | This option would raise the standard and scope of regulation without affecting trophy hunting where it is well-regulated and benefits species and people. Specifically, an import permit would be required for hunting trophies from species in Annex B of the WTRs. And imports of trophies from all species on Annexes A and B of the WTRs would be required to demonstrate conservation and other benefits (e.g., revenue sharing with local communities), which would be legally binding, to qualify for an import permit. This option would incentivize improved management and transparency of hunting operations and encourage reform of bad practice (e.g., canned hunting). The UK already makes NDFs at smaller spatial scales than species-country combinations (Fleming 2024). This policy option would not risk negatively impacting wildlife and/or local communities or Indigenous peoples that live in areas where trophy hunting takes place. | An assessment or certification system may be required for operators, which could be time-consuming to establish and develop at scale. |

## Footnotes

1 This excludes the Javan langur (*Trachypithecus auratus*) (2 live animals exported to South Africa in 2002) and the tiger (*Panthera tigris*) (1 pre-Convention [source code O] skin exported to Mexico in 2019).

## Notes

### Competing Interest Statement

DWSC is a member of the IUCN CEESP/SSC Sustainable Use and Livelihoods Specialist Group (SULi) and the IUCN SSC Pangolin Specialist Group.
MtSR is a member of the IUCN SSC African Rhino Specialist Group and SULi.
AD conducted this work under a Fellowship funded by the Recanati-Kaplan Foundation and Panthera. She has consultancies with the Darwin Expert Committee and Jamma International, but neither funded this work. AD leads WildCRU which has funding from donors with a wide variety of views of trophy hunting.
DH receives research funding from Jamma International, WWF Deutschland, and the Luc
Hoffmann Institute (now Unearthodox), the Band Foundation, and the John Muir Trust.
AH is a member of SULi and has received non-personal funding from Jamma International, although not for this study.
MH is employed by ZSL and is a member of the IUCN SSC Afrotheria, Antelope, Bear, and Canid Specialist Groups, and SULi.
RM-C is employed as the Chief Ecologist Terrestrial with the Parks and Wildlife Management Authority in Zimbabwe.
DR is the Chair of SULi which receives funding from Jamma International and the Abu Dhabi Environment Agency although neither funded this study, and is a member of the UK Advisory Group on Illegal Wildlife Trade and Darwin Expert Committee.

### Summary of Updates

This version incorporates edits to reflect the evolving policymaking processes in the UK and other edits based on a final review of the article by all authors.

